# A Minimal Synthetic Cleavage Module Enables Selective Degradation of Pathogenic r(CUG) Repeats in Cells

**DOI:** 10.64898/2026.11.04.764312

**Authors:** Jingsong Shan, Sandra Kovachka, Kisu Sung, Matthew D. Disney

## Abstract

Although direct RNA degradation offers an alternative to occupancy-driven activity, it has relied on complex natural product-derived cleavage modules such as bleomycin. Here, we show that a compact synthetic Fe(II)-coordinating warhead can be programmed for selective RNA degradation in cells. Coupling this cleavage module to a small molecule that recognizes expanded r(CUG) repeats that cause myotonic dystrophy type 1 (DM1) afforded selective degradation of the pathogenic RNA while sparing wild-type and transcripts harboring short r(CUG) repeats. In DM1 patient-derived myoblasts, degradation of mutant repeat RNA reduced DM1-associated pathology including formation of nuclear foci, sequestration of MBNL1, and alternative pre-mRNA splicing defects, without inducing DNA damage. These studies establish that minimal synthetic cleavage modules can replace complex natural product warheads to enable selective degradation of disease-causing RNAs in cells.

**Table of Contents:** 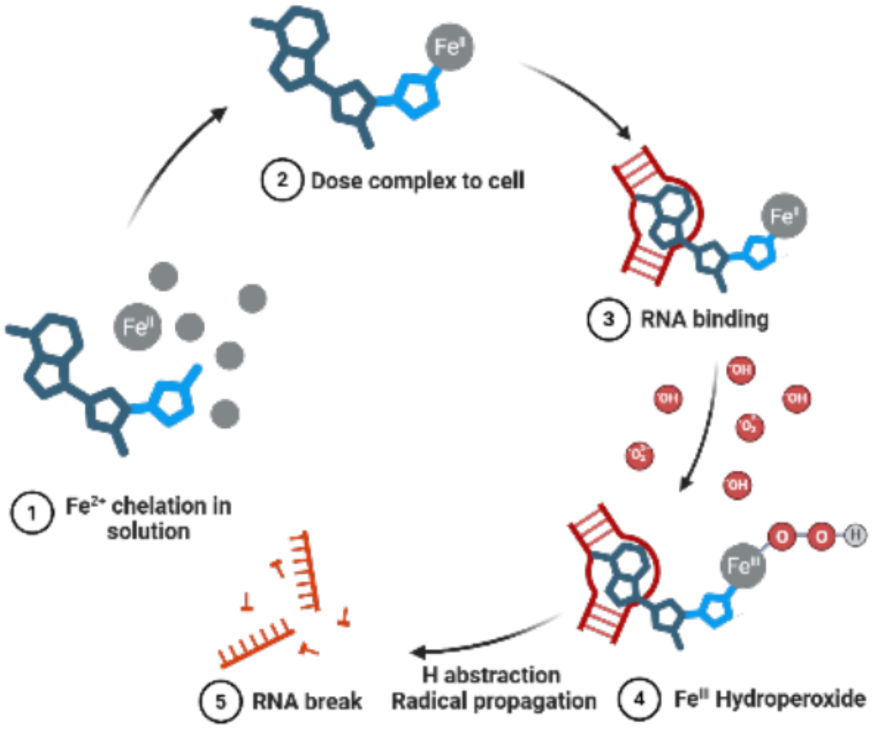

A compact synthetic Fe(II)-coordinating cleavage module is programmed to selectively degrade a pathogenic RNA target in cells. Coupling this warhead to an r(CUG)^exp^-binding small molecule enables targeted destruction of mutant DMPK transcripts and rescue of DM1-associated molecular phenotypes with no detectable DNA damage.

## Introduction

RNA structure plays central roles in cellular regulation by controlling translation, splicing, localization, and RNA stability [1–4]. Structured RNAs also contribute directly to human disease and therefore represent attractive therapeutic targets. Among these disorders, microsatellite repeat expansion diseases are particularly compelling because the pathogenic RNAs fold into repetitive structural motifs that sequester RNA-binding proteins (RBPs) [5–6]. Myotonic dystrophy type 1 (DM1) is caused by expansion of a CTG trinucleotide repeat within the 3′ untranslated region of the dystrophia myotonica protein kinase (DMPK) gene. Expanded alleles are transcribed into r(CUG)^exp^ RNA, which folds into a stable hairpin structure containing repeating 1×1 nucleotide U/U internal loops [7–8]. These motifs avidly bind the alternative pre-mRNA splicing regulator muscleblind-like 1 protein (MBNL1), resulting in sequestration of MBNL1 into nuclear foci and widespread dysregulation of splicing [9–13].

Small molecules that bind r(CUG)^exp^ can partially rescue MBNL1 function by competitively disrupting RNA–protein interactions [14–15]. However, reversible occupancy faces a major mechanistic limitation: pathogenic long repeat RNAs contain dozens to thousands of repeating structural motifs, requiring sufficient occupancy to achieve robust functional rescue [16]. In addition, non-covalent small molecules must bind r(CUG)^exp^ with very high affinity to efficiently compete with the avid MBNL1 protein [17]. In contrast, degraders can eliminate the entire pathogenic transcript following a single productive cleavage event, thereby circumventing the stoichiometric and affinity limitations of reversible binding. Previous studies demonstrated that direct cleavage of r(CUG)^exp^ can be achieved using bleomycin conjugates [18–19] or imidazole-based cleavage motifs [20–22]. While effective, bleomycin relies on a complex natural product scaffold associated with DNA cleavage activity and clinical toxicities [23], whereas the activity of imidazole requires a precise structural context and binder-driven destabilization of the RNA target [21]. We therefore sought to develop a modular and synthetically accessible direct degrader (unlike bleomycin) in which RNA recognition and cleavage activity are separable functional elements (unlike imidazole).

To accomplish this, we linked a high-affinity *bis*-benzimidazole dimer RNA-binding module [19] to a compact iron-coordinating oxidative cleavage warhead lacking intrinsic RNA affinity (1, Fig. 1A) [24–26]. We hypothesized that selective RNA binding would spatially position the cleavage module at the intended target, enabling direct degradation of r(CUG)^exp^ while minimizing off-target nucleic acid damage.

**Figure 1.**
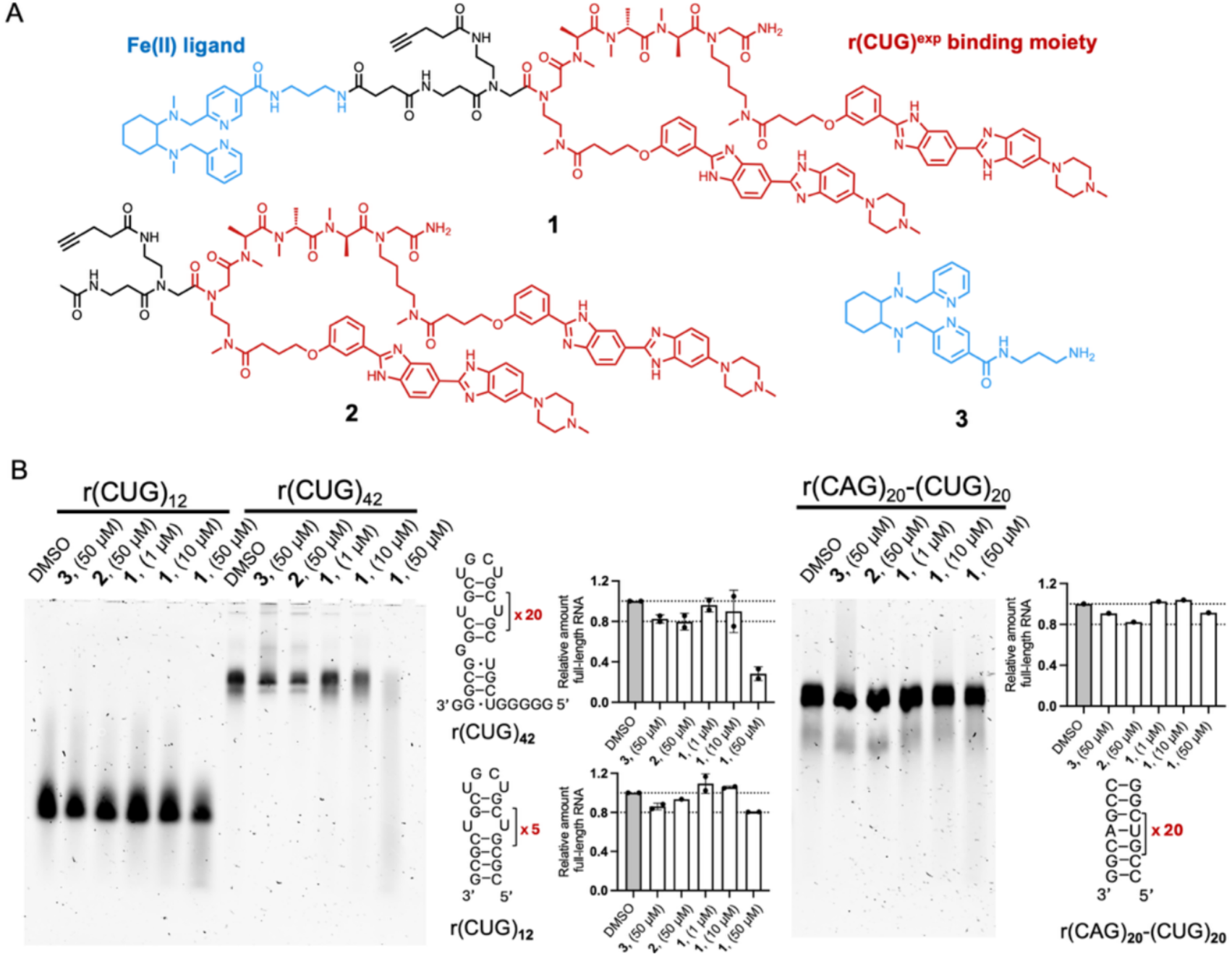
Heterobifunctional compound 3 selectively degrades r(CUG)^exp^ *in vitro*. (A) Chemical structures of heterobifunctional compound 1, r(CUG)^exp^ binder 2 and Fe(II) coordinating cleavage module 3. (B) In vitro cleavage assays showing selective degradation of r(CUG)^exp^ for 1 (right panel) and no significant cleavage for the binder (2) or the cleavage module (3) alone. A corresponding harboring 20 AU pairs instead of UU loops was completely resistant to cleavage by compound 1.

## Results and Discussion

### Design of a Modular Dimer Direct Degrader

To selectively recognize r(CUG)^exp^, we designed compound 1 by incorporating previously described *bis*-benzimidazole dimer 2 that binds repeating 1×1 nucleotide U/U internal loops with high affinity [19]. This RNA-binding scaffold was tethered to a tetradentate pyridine ligand 3 (Fig. 1A), a synthetic mimic of bleomycin, capable of generating localized oxidative cleavage chemistry [25, 27–28]. Upon Fe(II) chelation, the latter generates Fe(III)-OOH active species which in turn initiates H-abstraction and radical degradation of the backbone of nucleic acids [28–30]. Proven to induce efficient DNA cleavage, we evaluated the applicability of this approach for selective degradation of target RNAs. The resulting heterobifunctional molecule 1 (Fig. 1A) consists of two functionally distinct modules: (i) a selective RNA recognition element and (ii) an Fe(II) coordinating oxidative cleavage module lacking intrinsic RNA-binding affinity. We hypothesized that, rather than generalized cleavage of cellular nucleic acids, selective, productive cleavage would occur only after RNA binding positions the cleavage module in close proximity to the target RNA.

### The *Bis*-benzimidazole Dimer Direct Degrader Selectively Cleaves r(CUG)^exp^ *in vitro*

The activity of 1 was first evaluated using *in vitro* cleavage assays with r(CUG)_12_ and r(CUG)_42_ (Fig. 1B and S1A). RNA substrates were folded and incubated with compounds pre-complexed with an Fe(II) source and a reductant. The heterobifunctional degrader 1 efficiently cleaved both r(CUG)_12_ and r(CUG)_42_ RNAs within 2 h (Fig. 1B, left panel), with substantially stronger activity observed against the longer repeat construct, ∼80% cleavage for r(CUG)_42_ and ∼20% for r(CUG)_12_. Enhanced cleavage of r(CUG)_42_ likely reflects improved multivalent engagement of the *bis*-benzimidazole dimer along extended arrays of U/U internal loops and represent a more realistic proxy of the effect of the compound in the disease system where hundreds of r(CUG) repeats are present.

Importantly, neither the binder 2 alone nor the cleavage module 3 alone induced significant RNA degradation under identical conditions (Figs. 1B, left panel and S1A), demonstrating that both RNA recognition and oxidative cleavage functionalities are required. To assess selectivity, we evaluated cleavage against a base paired control RNA in which the U/U internal loops were replaced by canonical AU base pairs. These RNAs were completely resistant to cleavage even at concentrations that efficiently degraded r(CUG)^exp^ (Fig. 1B, right panel).

Further validation of the cleavage reaction for r(CUG)_42_ showed time- and dose-dependent decrease of full-length RNA, with degradation starting as soon as 30 min (∼20% cleavage at the highest 50 μM dose) and maximizing at 2 h reaction time (Fig. S1B). Addition of an oxidant increased the reaction rate for 1 (60% cleavage achieved by 5-10 μM of 1 after 30 min, Fig. S1C) along with non-specific cleavage given by the tetradentate cleavage module 3, suggesting a trickier fine-tuning required for selective RNA degradation under these conditions, with no distinct cleavage product (Fig. S1D).

Collectively, these data demonstrate that tetradentate iron coordinating warheads can induce selective RNA cleavage when tethered to selective RNA binders.

### The Degrader Selectively Reduces r(CUG)^exp^-containing Transcripts in Cells

To assess the cellular potency and selectivity of 1, HeLa480 cells that stably express a mutant interrupted r(CUG)_480_ allele composed of 24 tandem modules of [r(CUG)_20_(CUCGA)], and a wild type r(CUG)_0_ allele were employed [31]. Allele-specific TaqMan probes enable quantitative discrimination of the two transcripts by RT-qPCR. After 24h treatment, 1 dose dependently reduced the r(CUG)^exp^ transcripts with maximum effects observed at 10 μM (35 ± 3% reduction, p = 0.0002) without affecting wild type r(CUG)_0_ transcript level (Fig. 2A). Notably, neither the binder 2 nor warhead 3 alone altered RNA abundance, confirming that selective degradation requires the combined action of the RNA binder and direct degrader modules (Fig. 2A). To probe functional competition with MBNL1, we used a previously established live-cell NanoBRET assay that reports on MBNL1–r(CUG)^exp^ complex formation.[32] The degrader 1 dose dependently reduced NanoBRET signal with maximum inhibition rate of 72 ± 7% at 10 μM (p = 0.0003). In contrast, the dimeric binder 2 produced only a modest decrease in BRET with maximum reduction of 32 ± 6% at 10 μM (p = 0.0101), consistent with reversible displacement of MBNL1 rather than cleavage-driven elimination of the r(CUG)_480_ transcript (Fig. 2B). This enhancement confirms that coupling the binder to a degrader module that transforms a reversible inhibitor of MBNL1 binding into an RNA-cleaving agent leads to cellular effects unattainable with the binder alone.

**Figure 2.**
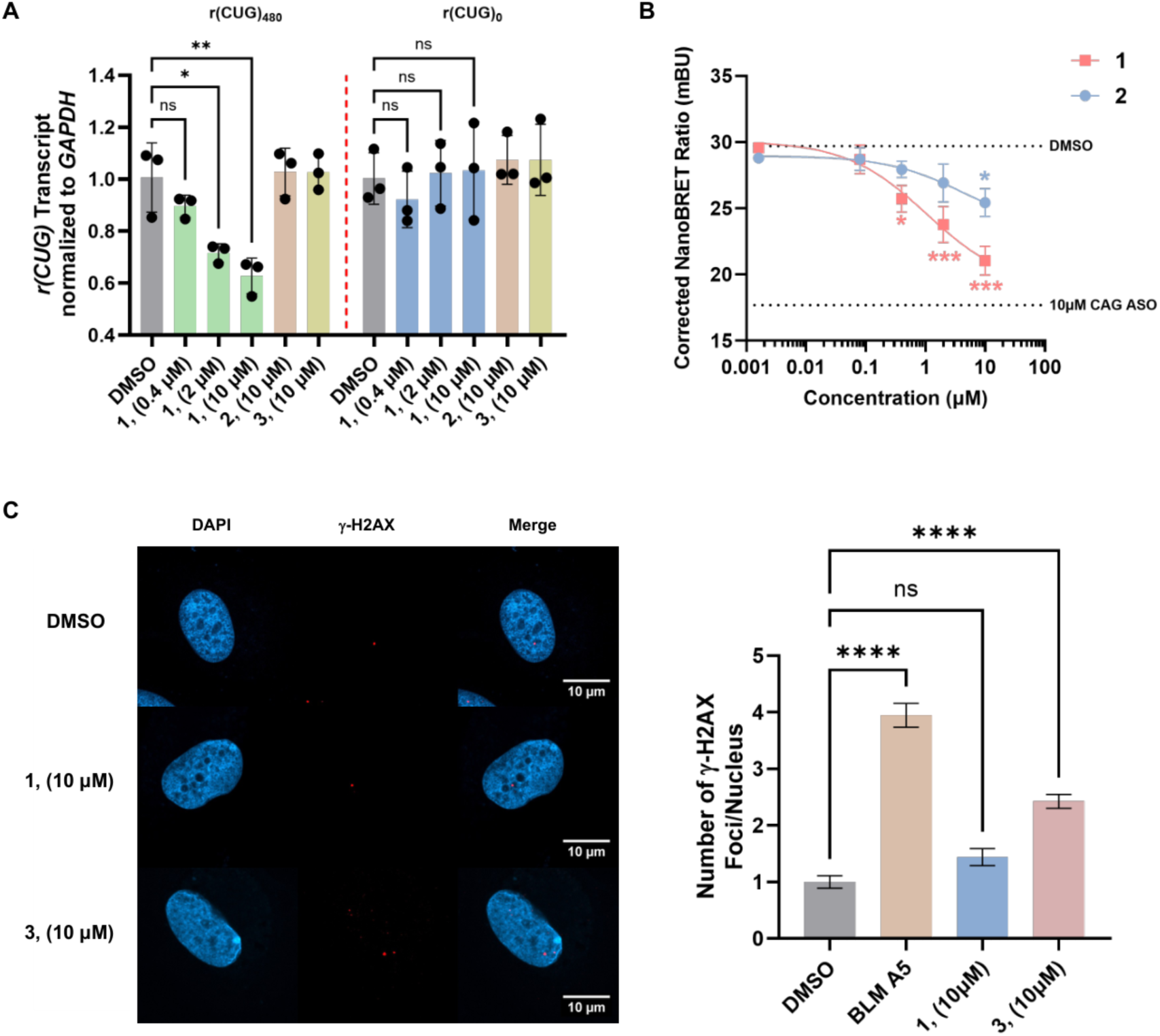
Cellular activity and selectivity of the bis-benzimidazole dimer direct degrader 3. (A) Effect of 1 – 3 on r(CUG)_480_ and wild-type DMPK transcripts in HeLa480 cells following treatment with the degrader 1 (0.4– 10 μM), as determined by RT-qPCR. Data are reported as mean ± standard deviation (n = 3 biological replicates). (B) NanoBRET analysis of MBNL1–r(CUG)^exp^ interactions in live cells following treatment with 1 and 2 (0.0016– 10 μM, n = 3 biological replicates). (C) Left: Representative images for γ-H2AX foci (red) and DAPI staining (blue) in HeLa480 cells treated with 1 or 3 to assess DNA damage (n = 3 biological replicates; 40 nuclei per replicate). Right: Quantification of γ-H2AX foci from imaging studies. For all panels, statistical significance was determined by a One-way ANOVA with multiple comparisons with significance thresholds: *, p < 0.05; **, p < 0.01; ***, p < 0.001; and ****, p < 0.0001.

### No Measurable DNA Damage Induced by the Degrader

Because oxidative cleavage chemistry can potentially damage DNA, we quantified phosphorylated H2A variant H2AX (γ-H2AX), a canonical marker of DNA double-strand breaks, to assess whether 1 elicits any DNA-associated off-target effects in cells [33]. In HeLa480 cells, a fluorescence resonance energy transfer (FRET) based assay foci with antibody targeting phosphorylated motif and H2AX respectively, was employed to quantify γ-H2AX after treating with the compound of interest for 24h (Fig. S2A). A cell-lysate control was incorporated to establish the assay window, yielding an approximate 2.8-fold range. The positive control BLM A5 produced a robust increase in γ-H2AX FRET signal at 10 µM (1.9 ± 0.1-fold, p < 0.0001), in agreement with previously published observations [18]. The direct module 3 alone also induce 1.5 ± 0.1-fold (p < 0.0001) signal increase at 10 µM, consistent with its previously reported potential to generate double-strand DNA breaks [26]. In contrast, treatment with the direct degrader 1 produced no detectable increase in γ-H2AX FRET signal up to 10 µM, indicating it does not elicit DNA-associated off-target effects and remains selective for its RNA target (Fig. S2B). To provide orthogonal validation, γ-H2AX immunostaining was performed to quantify nuclear foci. Both bleomycin A5 (p <0.0001) and 3 (p <0.0001) induced a significant increase in γ-H2AX foci per nucleus, whereas 1 (p = 0.12) had no measurable effect. Moreover, 1 displayed substantially reduced cytotoxicity compared to bleomycin-conjugated version 4. At 20 µM, 1 preserved >90% cell viability, whereas 4 reduced viability to 74 ± 4%. This divergence further widened at 50 µM, where 1 maintained > 75% viability while 4 decreased cell viability to 37 ± 4% (Fig. S3). These results demonstrate that appending the RNA-binding modules reprograms bleomycin’s cleavage selectivity, restricting its activity to the intended RNA target in cells, thereby minimizing off-target DNA cleavage and cellular toxicity.

### Correlating Degradation with Rescue of pre-mRNA Splicing Defects and Formation of Foci

The bioactivity of 1 was next evaluated in DM1 patient-derived and healthy myoblasts. For these studies, DM1 patient-derived or wild-type fibroblasts were forced into myogenic differentiation by doxycycline-induced expression of myoblast determination protein 1 (MYOD1) [34]. The patient-derived myoblasts express mutant (2600 r(CUG) repeats) and wild type alleles. Following 24 h of MYOD1-induced differentiation, the cells were treated with compounds in differentiation medium for a further 24 h. The ability of 1 to cleave r(CUG)_2600_ in the 3′ UTR of DMPK was then measured by reverse transcription-quantitative polymerase chain reaction (RT-qPCR) and droplet digital (dd)PCR. The small molecule dose dependently cleaved total DMPK mRNA with maximum of 38 ± 4% (p = 0.0017) at 10 µM without effects in WT myoblast (Fig 3A). To determine whether this selectivity extends to discrimination between expanded and short r(CUG) repeats, we next examined the effect of 1 on five endogenous transcripts containing short r(CUG) tracts with repeat lengths of 7-16. In contrast to the observed reduction of DMPK abundance, none of these RNAs were affected by treatment with 10 µM of degrader 1 (Fig. 3B). Collectively, these data demonstrate that 1 cleaves RNA in a repeat-length dependent manner, preferentially targeting expanded r(CUG) tracts.

**Figure 3.**
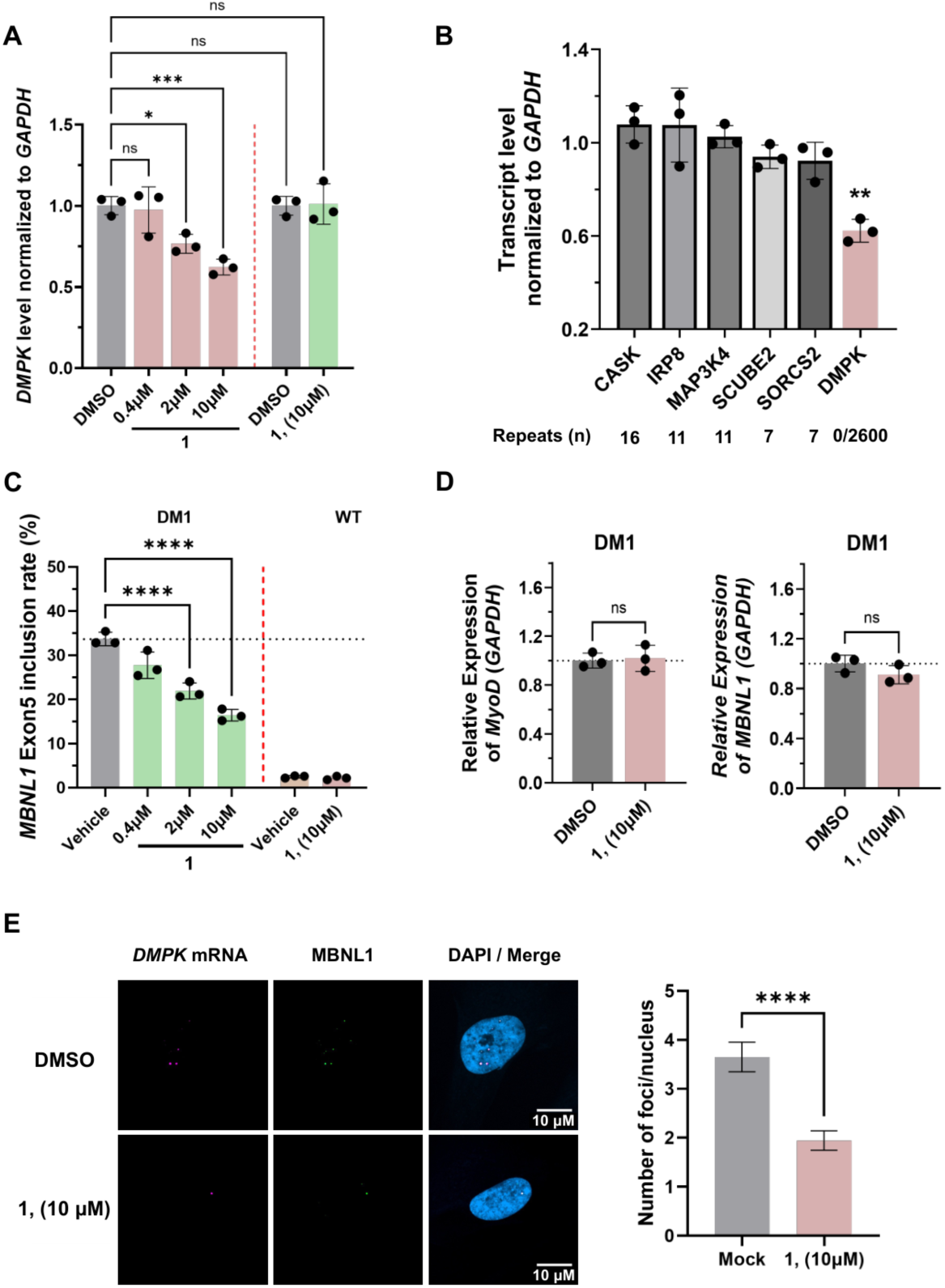
Rescue of DM1 molecular phenotypes in patient-derived myoblasts. (A) RT–qPCR analysis of DMPK transcripts in DM1 myoblasts (red) following treatment with the degrader 1 (0.4– 10 μM) and in WT myoblast treated with 10 μM of 1 (green). Data are reported as mean ± standard deviation (n = 3). (B) Effect of 1 (10 μM) on endogenous transcripts containing short r(CUG) tracts with repeat lengths of 7-16, in comparison to DMPK transcripts, in DM1 myoblasts, as determined by RT-qPCR (n = 3 biological replicates). Statistical significance was determined relative to vehicle (0.1% (v/v) DMSO) using a two-tailed unpaired Student’s t test with significance thresholds: *, p < 0.05; **, p < 0.01; ***, p < 0.001. (C) Effect of 1 on MBNL1 exon 5 alternative splicing in DM1 and WT myoblasts, as measured by end-point RT-PCR. (n = 3 biological replicates) Statistical significance was determined by a One-way ANOVA with multiple comparisons with significance thresholds: *, p < 0.05; **, p < 0.01; ***, p < 0.001; and ****, p < 0.0001. (D) Effect of 1 (10 µM) on MyoD and MBNL1 transcript abundance in DM1 myoblasts, as determined by RT-qPCR (n = 3 biological replicates). (E) Single-molecule RNA fluorescence *in situ* hybridization (smRNA-FISH) for DMPK CDS (magenta) and immunohistochemistry for MBNL1 (green). Left: Representative images of DM1 patient-derived myoblasts treated with 10 µM of 1 or vehicle (0.1% (v/v) DMSO). DAPI staining (blue) was used to visualize nuclei. Scale bar is 10 µm. Right: Quantification of r(CUG)^exp^-MBNL1 foci in the nucleus of DM1 myoblasts (40 nuclei quantified/replicate; n = 3 biological replicates). ****, p < 0.0001, as determined by Student’s t-test. Data are reported as the mean ± SEM.

The activity of 1 was then evaluated for its ability to improve DM1-associated cellular phenotypes in patient-derived myoblasts, specifically the rescue of alternative pre-mRNA splicing defects and the reduction of nuclear RNA foci [35–37]. After 24h myogenic differentiation, cells were treated with the compound of interest during myogenic differentiation for another 24h, followed by measuring rescue of the MBNL1 exon 5 alternative splicing defect observed in this cell line [34]. In the absence of compound treatment, the rate of exon 5 inclusion in DM1 patient-derived myoblasts was 34 ± 2%, while the rate of inclusion in differentiated wild type myoblast was 4 ± 1% (Fig. 3C). Compound rescue levels were calculated as the fractional decrease in exon 5 inclusion rate relative to untreated DM1 myoblasts, normalized to the DM1 value (set to 100%) and the wild-type value (set to 0). The degrader 1 dose dependently rescued MBNL1 exon 5 alternative splicing with the most potent effects at 10 µM reducing exon 5 inclusion rate from 34 ± 2% to 16 ± 1% (54 ± 4% rescue, p <0.0001) (Figs. 3C and S4). An additional MBNL1-regulated splicing event, SOS1 exon 25, was also improved dose dependently upon treatment with 1 (Fig. S5) [38]. Notably, 1 had no effect on MBNL1 exon 5 splicing in wild-type myoblasts, nor did it alter MBNL1-independent splicing events including MAP4K4 exon 22a and ANK2 exon 21 in DM1 myoblasts (Fig. S6) [38–40]. Additional qPCR analysis confirmed that the transcript abundances of MBNL1 and MyoD1 were not affected by 10 µM of 1 under the same conditions to evaluate alternative splicing, confirming the observed splicing rescue does not arise from off-target effects (Fig. 3D).

To assess whether the small molecules alleviate MBNL1 sequestration by r(CUG)^exp^, the number of MBNL1-positive, r(CUG)^exp^ nuclear foci in DM1 myoblasts was quantified using immunofluorescence (IFA) together with single molecule fluorescence *in situ* hybridization (smFISH) probes targeting DMPK exons 11–15, as previously described [32]. Treatment with 10 µM of 1 significantly reduced the number of r(CUG)^exp^−MBNL1 nuclear foci (1.8 ± 0.1 RNA foci/nuclei, p < 0.0001) in DM1 myoblasts compared to vehicle-treated samples (3.4 ± 0.1 RNA foci/nuclei) (Fig. 3E). This reduction is consistent with compound-induced degradation of the pathogenic transcript and release of sequestered MBNL1.

### Transcriptome-wide Analysis Demonstrates 1 Improves DM1-associated Defects Broadly

To comprehensively assess the effects of 1 in patient-derived DM1 myoblasts, transcriptome-wide analysis using Nanopore long-read sequencing was performed on total RNA isolated from differentiated DM1 and WT myoblasts. Consistent with prior RT-qPCR measurements (Fig. 3A), RNA-seq analysis of genes containing short, nonpathogenic r(CUG) repeats expressed in DM1 myoblasts did not significantly alter after treatment of 10μM 1 (Fig. S7). However, because the mutant DMPK allele contains ∼2600 r(CUG) repeats, the expanded transcript was lost during cDNA-PCR library preparation, which is prone to polymerase slippage and allele dropout, a phenomenon well documented in trinucleotide repeat disorders [41–42]. In contrast, the WT allele was retained, and no reduction in its abundance was observed, confirming the specificity of the direct degrader 1. We next investigated the effect of 1 on 56 transcripts encoding proteins involved in the DNA-damage pathway [43]. These transcripts remained unchanged in DM1 myoblast following treatment with 10 μM of 1 (Fig. S7), in agreement with γ-H2AX foci studies (Fig. 2C). As expected, expression of r(CUG)^exp^ caused extensive transcriptome-wide dysregulation in DM1 myoblasts. Comparison of untreated DM1 and WT myotubes identified 728 genes with significantly altered abundance (baseMean > 10, log2 fold enrichment > 0.5 and false discovery rate (FDR) < 0.05). Upon treatment with 1 (10 μM), 322 deregulated genes (48%) were shifted toward WT levels, where their abundances in treated DM1 myoblasts and untreated WT myoblasts were no longer statistically different (FDR > 0.05) (Fig. 4A). Moreover, consistent with qPCR measurements of endogenous transcripts containing short, non-pathogenic r(CUG) tracts (Fig. 3B), expression of CASK and other short-repeat–containing genes remained unaffected by 1-treatment (Fig. S8A).

**Figure 4.**
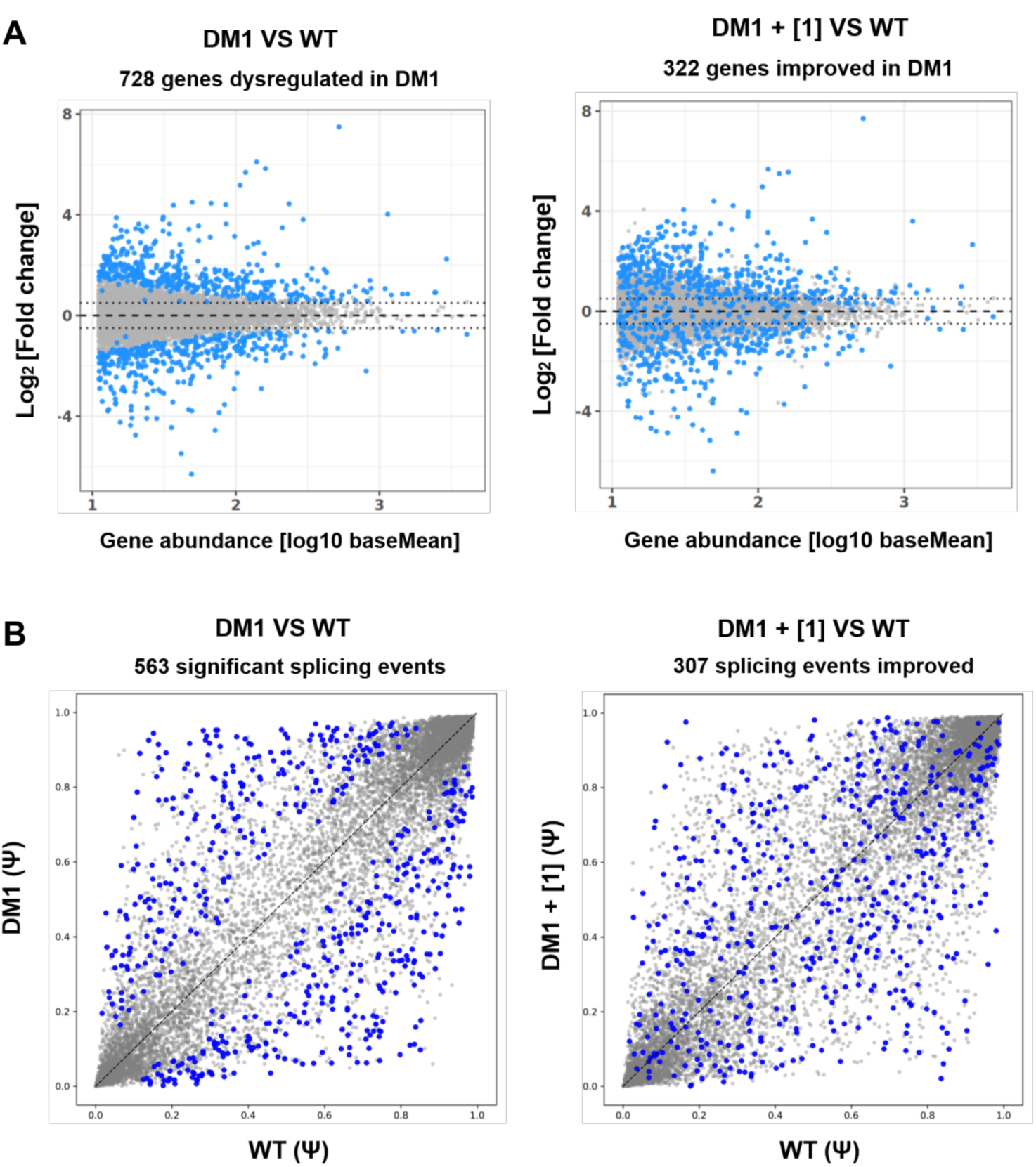
Transcriptome-wide assessment of compound 1 in DM1 patient-derived myoblasts. (A) Left: Differential gene-expression analysis comparing DMSO-treated DM1 patient-derived myoblasts with DMSO-treated WT myoblasts. Genes significantly dysregulated in DM1 myoblasts (baseMean > 10, log2 fold enrichment > 0.5 and FDR < 0.05; n = 2) are highlighted in blue. Right: Differential gene-expression analysis of DM1 myoblasts treated with 10 μM of 1 compared to DMSO-treated WT myoblasts. Genes highlighted in blue are significantly dysregulated in DM1 myoblasts ((|ΔΨ| >0.1, FDR < 0.05; n = 2). (B) Left: Alternative splicing events in DMSO-treated DM1 vs WT myoblasts. Right: Splicing analysis of DM1 myoblasts treated with 10 μM of 1 vs. DMSO-treated WT myoblasts. Blue points indicate significantly mis-spliced events in DM1 myoblasts (FDR < 0.05, n = 2). Events shifting toward the diagonal reflect rescue of splicing defects upon treatment with 1.

Given that sequestration of MBNL1 by r(CUG)^exp^ alters alternative splicing of its canonical substrates [9–13], we next analyzed the RNA-seq data for rescue of transcriptome-wide splicing events by 1. Transcriptome-wide analysis by SUPPA2 [44] revealed 563 significantly misregulated splicing events (|ΔΨ| >0.1 and FDR < 0.05) in untreated DM1 myoblasts as compared to WT myoblasts. Treatment with 10 μM of 1 restored 307 of these events (55%) toward WT splicing patterns, eliminating statistical differences (FDR > 0.05) (Fig. 4B). Notably, seven splicing events previously reported to correlate with DM1 molecular pathology and clinical manifestations [45–46] were also rescued by treatment with 10 μM of 1 (Fig. S8B). Collectively, these results demonstrate that conjugating a small molecule with cleavage module can confer potent and selective cleavage of an RNA target, rescuing downstream disease pathways in patient-derived muscle cells.

## Conclusion

The studies presented here establish a modular strategy for direct chemical degradation of structured RNAs. By coupling a selective a *bis*-benzimidazole dimer RNA-binding module to a minimized tetradentate Fe (II) chelating oxidative cleavage warhead, we converted reversible RNA binding into selective transcript destruction.

Several mechanistic features distinguish this degrader design. First, cleavage activity emerges only when RNA recognition and oxidative chemistry are physically linked. Neither module alone is sufficient to induce degradation. Second, the degrader preferentially targets expanded r(CUG) repeat tracts, highlighting the importance of multivalent engagement across the expanded structural motifs. Third, selective RNA targeting spatially constrains oxidative chemistry and prevents measurable DNA damage in cells despite the intrinsic reactivity of the free cleavage module. Furthermore, proximity-induced transcript degradation showed stronger MBNL1 displacement by ∼35% compared to reversible binding.

## Supporting Information

The authors have cited additional references within the Supporting Information [47–55]. In vitro cleavage assays were performed by gel electrophoresis with r(CUG)_12_ and r(CUG)_42_. Cellular activity was assessed using NanoBRET, cell viability assays, transcript level measurement by RT-qPCR and ddPCR, imaging of MBNL1-r(CUG)^exp^ foci as well as alternative pre-mRNA splicing analysis by end-point RT-qPCR analysis. DNA damage was evaluated by γ-H2AX TR-FRET and foci imaging. Complete experimental methods and materials, including synthetic methods and compound characterization, are provided in the Supporting Information.

## Acknowledgements

This work was funded by the U.S. National Institutes of Health R35 NS116846 (to M.D.D.), the Department of Defense (HT94252310336 to M.D.D.) and the Muscular Dystrophy Association (1069959 to M.D.D). We thank Dr. Nicolai Urban and Dr. Long Yan at Max Plank Florida Institute offering the LSM 880 microscope and imaging advice, Dr. Hugo Helbert for the synthesis of 3, and Dr. Álvaro Simba-Lahuasi for *in vitro* transcription of r(CUG)_42_.

