## Supplementary Information for "A Minimal Synthetic Cleavage Module Enables Selective Degradation of Pathogenic r(CUG) Repeats in Cells"

#### Supplementary Tables & Figures

**Table S1**. Sequences of primers used in these studies.

| | | |
|---|---|---|
| DNA templates used for *in vitro* transcription | | |
| ssDNA for r(CAG)20 | 5'-mGGCTGCTGCTGCTGCTGCTGCTGCTGCTGCTGCTGCTGCTGCTGCTGCT GCTGCTGCTGCTGCCTATAGTGAGTCGTATTA-3' (T7 promoter) | |
| ssDNA for r(CUG)20 | 5'-mGGCAGCAGCAGCAGCAGCAGCAGCAGCAGCAGCAGCAGCAGCAGCAG CAGCAGCAGCAGCAGCCTATAGTGAGTCGTATTA-3´ (T7 promoter) | |
| Primers and probes used for RT-qPCR amplification of HeLa cells stably expressing mutant and WT alleles (HeLa480)[1] | | |
| **Primers** | **Forward Sequence (5'→ 3')** | **Reverse Sequence (5'→ 3')** |
| HeLa *DMPK* WT and mutant alleles | CGATCTCTGCCTGCTTACTC | GTCGGAGGACGAGGTCAATAAA |
| *GAPDH* (SYBR) | GAAGGTGAAGGTCGGAGTC | GAAGATGGTGATGGGATTTC |
| **Probe** | **Sequence (5'→ 3')** | |
| Probe 1 (WT allele) | /56FAM/AGAGCAGCG/ZEN/CAAGTGAGGAGG/3IABkFQ/ | |
| Probe 2 (mutant allele) | /5HEX/TGACGCAGC/ZEN/CACGTGAAGGTC/3IABkFQ/ | |
| Primers used in MyoD-inducible WT and DM1 myoblasts [2-3] | | |
| | **Forward Sequence (5'→ 3')** | **Reverse Sequence (5'→ 3')** |
| *DMPK* | CGTGCAAGCGCCCAG | CTCCACCAACTTACTGTTTCATCCT |
| *MBNL1* exon 5 splicing | GCTGCCCAATACCAGGTCAAC | TGGTGGGAGAAATGCTGTATGC |
| *SOS1* exon25 splicing | TGGTGCTTCCAGTACCACAG | GGCAGATTCTGGTCGTCTTC |
| *MyoD* | CGGCATGATGGACTACAGCG | CAGGCAGTCTAGGCTCGAC |
| *CASK* | TTGAAATCGTAAAGCGAGCTGA | CAGTAGCGTAGAGCTTCCAGTA |
| *IRP8* | GCCAAGGATTGCGAAAAGGAC | GTGGTCTAAGCAGTCATCGTC |
| *MAP3K4* | CAATAAGCCTTACCTCAGCCTTG | GTTAAGCCAGAAACCAGACGTA |
| *SCUBE2* | CCCACCTCCTACAAGTGCTC | TGCAACGATAATTGCCTGGAAT |
| *SORCS2* | CACGTCGTTCGTGCTCAAG | CGTCCCGAAATCTGATGACCG |

| **Table S2.** Sequence of smFISH probe targeting *DMPK* used for foci imaging in DM1 myoblasts (previously reported) labelled with ATTO-550 on 3’ end [4]. | | |
|---|---|---|
| **Probe ID** | **Sequence (5’ → 3’)** | **Targeting nucleotide sequence (5’ → 3’) and nt range in *DMPK*** |
| DMPK_CDS_1 | AAGAAGTCGGCCACGTACTT | AAGTACGTGGCCGACTTCTT; 271-290 |
| DMPK_CDS_2 | CTTAAGCCTCACCACGATGG | CCATCGTGGTGAGGCTTAAG; 2628-2647 |
| DMPK_CDS_3 | GATCACCTTCAGAATCTCGA | TCGAGATTCTGAAGGTGATC; 2673-2692 |
| DMPK_CDS_4 | GTCTGCTTCATCTTCACTAC | GTAGTGAAGATGAAGCAGAC; 2720-2739 |
| DMPK_CDS_5 | GTTCATGATCTTCATGGCAT | ATGCCATGAAGATCATGAAC; 2751-2770 |
| DMPK_CDS_6 | TCACGGAAGCACGACACCTC | GAGGTGTCGTGCTTCCGTGA; 2795-2814 |
| DMPK_CDS_7 | CGAAGTGCAGCTGCGTGATC | GATCACGCAGCTGCACTTCG; 2851-2870 |
| DMPK_CDS_8 | TACAGGTAGTTCTCATCCTG | CAGGATGAGAACTACCTGTA; 2876-2895 |
| DMPK_CDS_9 | CACGTAATACTCCATGACCA | TGGTCATGGAGTATTACGTG; 2898-2917 |
| DMPK_CDS_10 | AAACTTGCTCAGCAGTGTCA | TGACACTGCTGAGCAAGTTT; 2931-2950 |
| DMPK_CDS_11 | AGTCTATGGCCATGACAATC | GATTGTCATGGCCATAGACT; 2995-3014 |
| DMPK_CDS_12 | ATGTTGTCGGGTTTGATGTC | GACATCAAACCCGACAACAT; 3044-3063 |
| DMPK_CDS_13 | TTGAGGCAAGAGCCGAAGTC | GACTTCGGCTCTTGCCTCAA; 3098-3117 |
| DMPK_CDS_14 | CTGCCCATAGAACATTTCAT | ATGAAATGTTCTATGGGCAG; 3267-3286 |
| DMPK_CDS_15 | CGTGGAATCCGCGTAGAAGG | CCTTCTACGCGGATTCCACG; 3291-3310 |
| DMPK_CDS_16 | ACGATCTTGCCATAGGTCTC | GAGACCTATGGCAAGATCGT; 3314-3333 |
| DMPK_CDS_17 | GAGAGAGGTGCTCCTTGTAG | CTACAAGGAGCACCTCTCTC; 3337-3356 |
| DMPK_CDS_18 | AGCAACCGCTGAATGAAGTC | GACTTCATTCAGCGGTTGCT; 3395-3414 |
| DMPK_CDS_19 | AAGAAGGGATGTGTCCGGAA | TTCCGGACACATCCCTTCTT; 3458-3477 |
| DMPK_CDS_20 | GAAATCCGGTGTAAAGGGGG | CCCCCTTTACACCGGATTTC; 3516-3535 |
| DMPK_CDS_21 | CACCAAGTCGAAGTTGCATG | CATGCAACTTCGACTTGGTG; 3552-3571 |
| DMPK_CDS_22 | CGAATGTCCGACAGTGTCTC | GAGACACTGTCGGACATTCG; 3608-3627 |
| DMPK_CDS_23 | CATGCAGGAGTAGGAGTAGC | GCTACTCCTACTCCTGCATG; 3666-3685 |
| DMPK_CDS_24 | ACTTCAGCTGTTTCATCCTG | CAGGATGAAACAGCTGAAGT; 3794-3813 |
| DMPK_CDS_25 | TAGTTGACTGGCGAAGTTCT | AGAACTTCGCCAGTCAACTA; 3951-3970 |
| DMPK_CDS_26 | CATCTAGATGGGAAGGTGGA | TCCACCTTCCCATCTAGATG; 4096-4115 |
| DMPK_CDS_27 | GAACAGGAGCAGGGAAAGCG | CGCTTTCCCTGCTCCTGTTC; 4224-4243 |
| DMPK_CDS_28 | CGGCACGAGACAGAACAACG | CGTTGTTCTGTCTCGTGCCG; 4246-4265 |
| DMPK_CDS_29 | CGGAGTCGAAGACAGTTCTA | TAGAACTGTCTTCGACTCCG; 4356-4375 |
| DMPK_CDS_30 | CTCGGAGCGGTTGTGAACTG | CAGTTCACAACCGCTCCGAG; 4436-4455 |
| DMPK_CDS_31 | GATCACAGGACTGGAGCTGG | CCAGCTCCAGTCCTGTGATC; 4469-4488 |

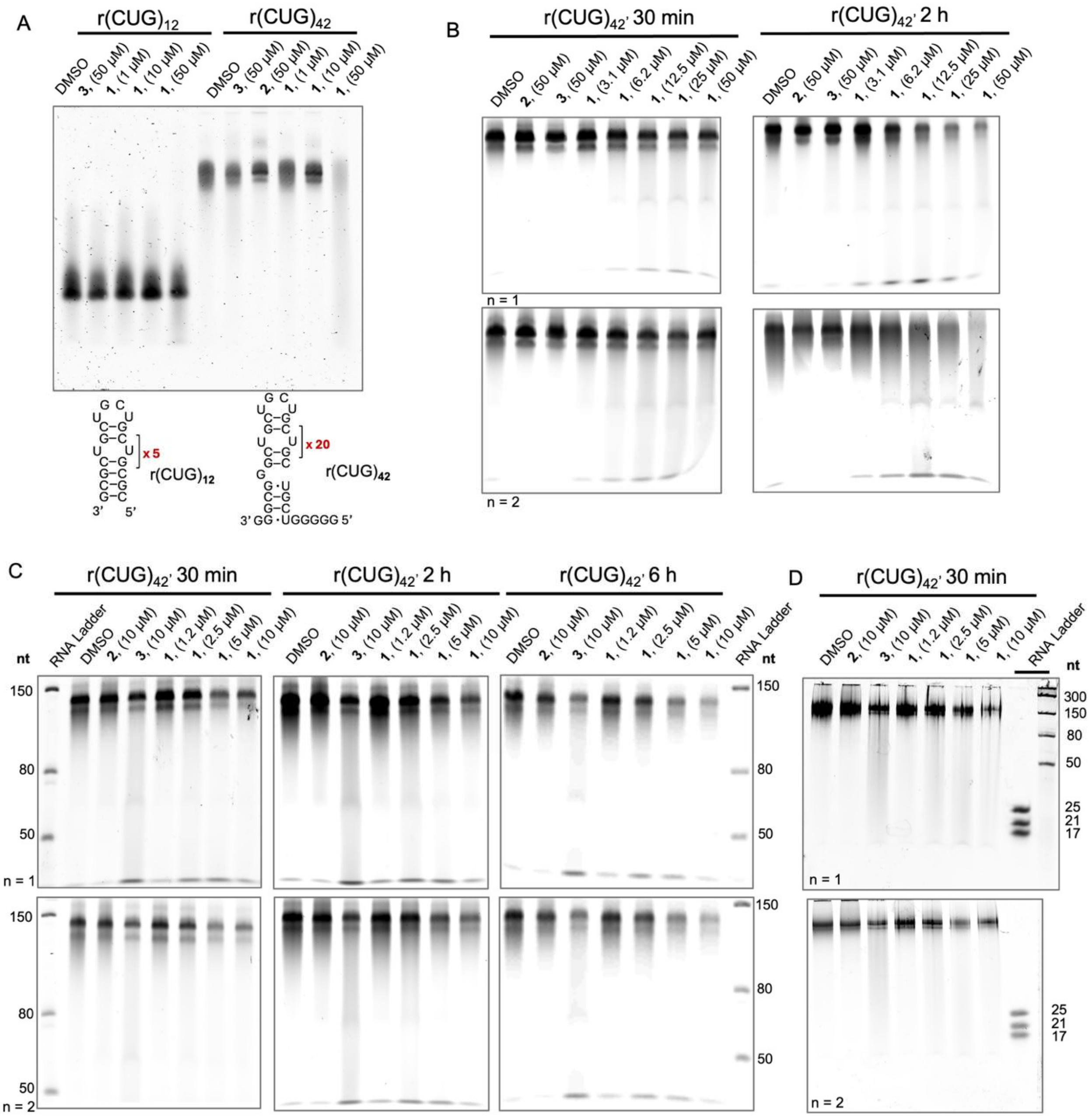

**Fig. S1.** Heterobifunctional compound **3** selectively degrades r(CUG)$^{exp}$ *in vitro*. (**A**) *In vitro* cleavage assay showing selective degradation of r(CUG)$^{exp}$ by **1** and no significant cleavage for the binder (**2**) or the cleavage module (**3**) alone. (**B**) Time and dose-dependent cleavage of r(CUG)$_{42}$ with maximum cleavage occurring after a 2 h incubation time. (**C**) Time and dose-dependent cleavage of r(CUG)$_{42}$ in the presence of oxidant (hydrogen peroxide). (**D**) No visible cleavage products could be identified via gel electrophoresis analyses (denaturing 20% (w/v) acrylamide gel, post-stained with SYBR Gold). For panels A-C, electrophoretic analyses was completed using denaturing 8% (w/v) polyacrylamide gels, and bands were visualized via post-staining the gel with SYBR Gold.

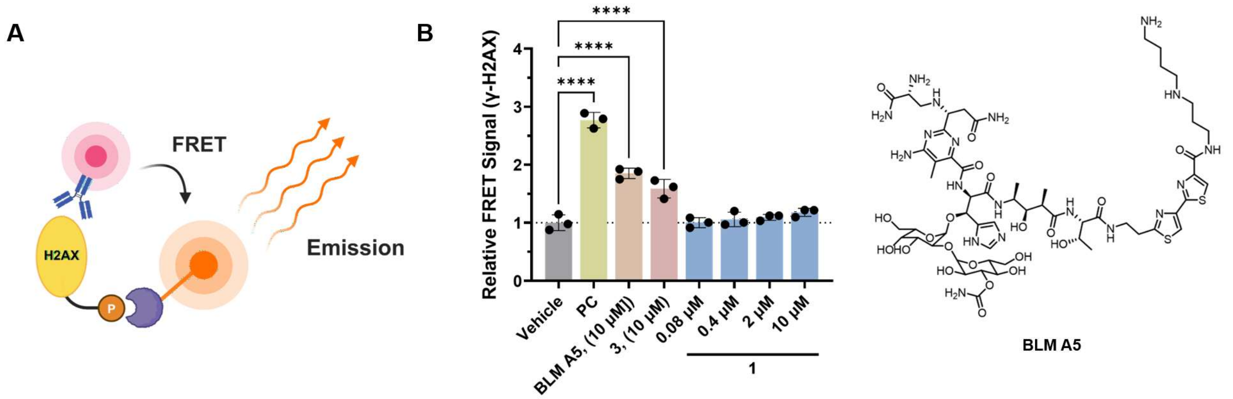

**Fig. S2. Effects of heterobifunctional 1 on γ-H2AX, a marker of DNA damage [5], in HeLa480 cell lysate.** (**A**) Schematic for measuring γ-H2AX in cell lysates using a time-resolved fluorescence resonance energy transfer (TR-FRET) assay. (**B**) Dose-dependent effects of **1** (0.4–10 μM), **3** (10 μM), and the natural product bleomycin A5 (BLM A5; 10 μM; structure shown on right) on γ-H2AX expression; and a positive control lysate (PC) was included to validate assay performance. Data are reported as mean ± standard deviation (n = 3 biological replicates), (**) P < 0.01, (***) P < 0.001, as determined by comparison to vehicle-treated cells (0.1% (v/v) DMSO) by a one-way analysis of variance (ANOVA).

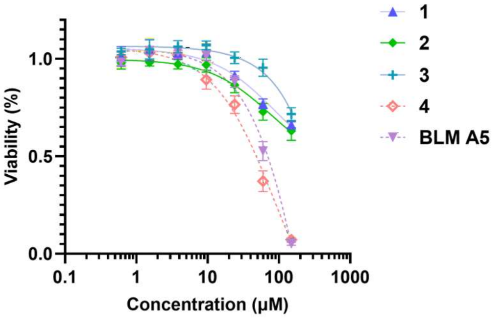

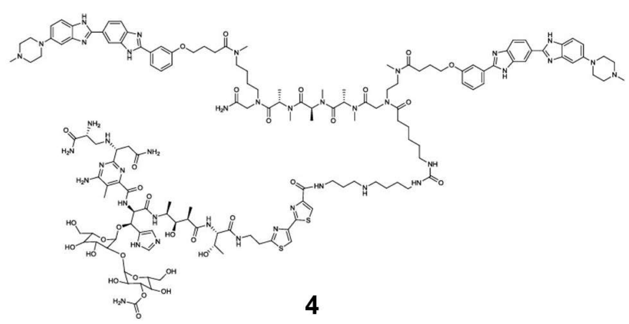

**4**

**Fig. S3. Effects of 1 – 3, bleomycin-conjugated derivative 4, and bleomycin A5 (BLM A5) on the viability of HeLa480.** Left: plot of the effect of the compounds on cell viability as a function of concentration (**BLM A5** structure shown in Fig. S2). Data are reported as the mean ± SD (n = 3 biological replicates). Right: chemical structures of **4**.

A

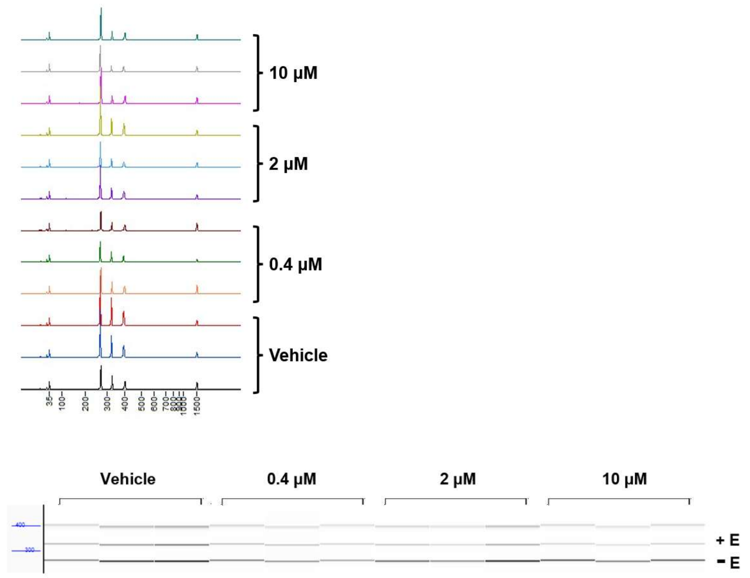

B

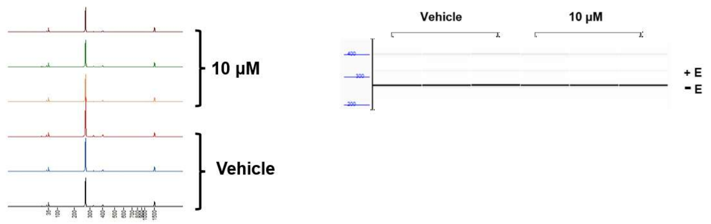

**Fig. S4. Effect of 1 on the alternative splicing of *MBNL1* exon 5 in DM1 patient-derived myoblasts.** (**A**) Representative electropherogram and pseudo-gel to study the effect of **1** on *MBNL1* exon 5 splicing, as assessed by end-point RT-PCR and fragment analyzer. (**B**) Representative electropherogram and pseudo-gel to study the effect of 10 µM of **1** on *MBNL1* exon 5 splicing in WT myoblasts, as assessed by end-point RT-PCR and fragment analyzer. In both panels, exon 5 inclusion is indicated by “+E” while exclusion is indicated by “-E” (n = 3 biological replicates). The additional band above “+E” represents a hybrid duplex formed by annealing between the exon5–containing and exon5–skipped amplicons and was excluded from quantification. The quantification of exon 5 inclusion can be found in Fig. 3C.

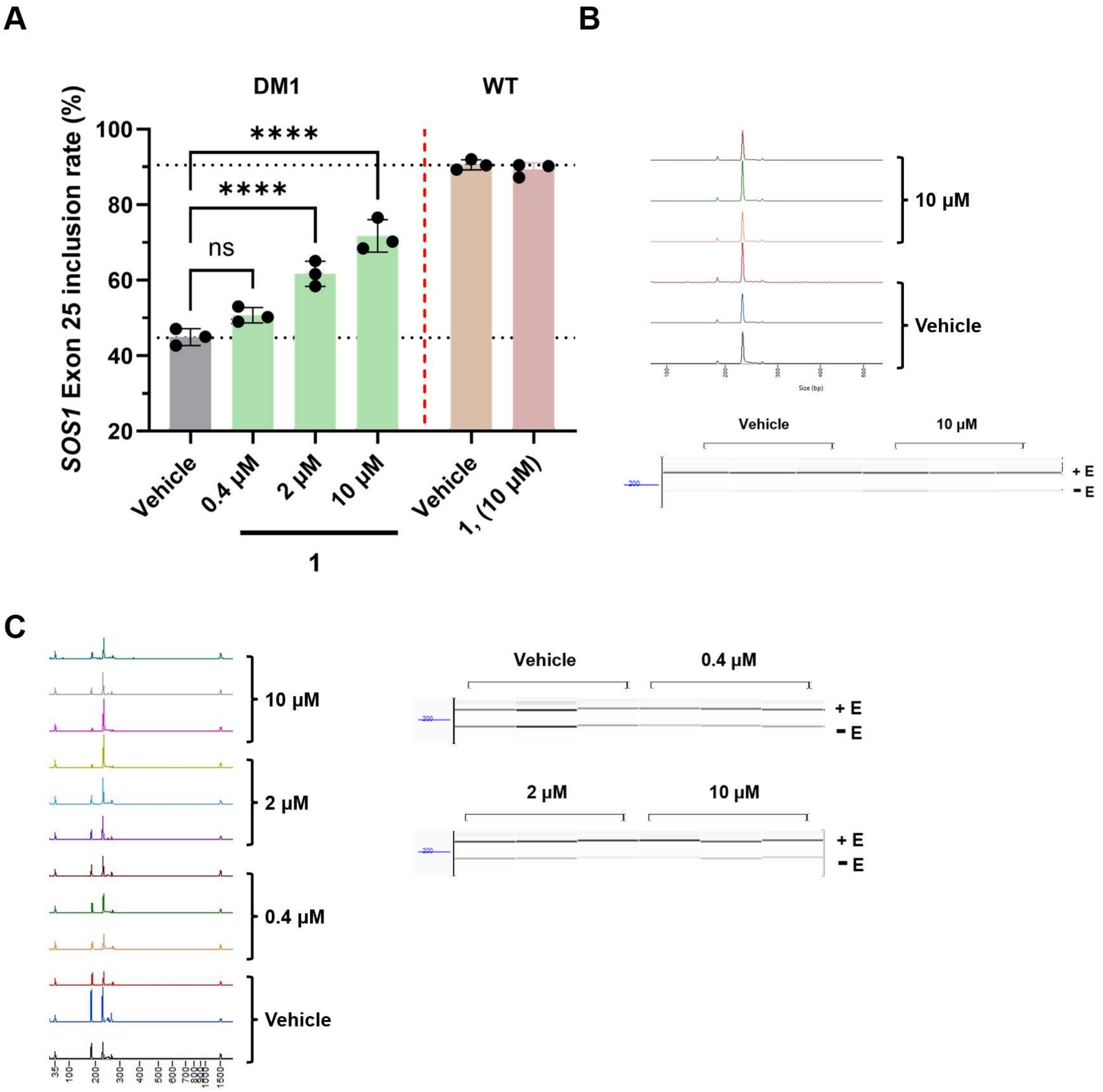

**Fig. S5. Fragment analyzer gel of *SOS1* exon 25 in myoblasts.** (**A**) Effect of **1** on *SOS1* exon 25 splicing in DM1 patient-derived and WT myoblasts, as assessed by end-point RT-PCR and fragment analyzer (n = 3 biological replicates). (**) P < 0.01, (***) P < 0.001, as determined by comparison to vehicle-treated myoblasts (0.1% (v/v) DMSO) by a one-way ANOVA. (**B**) Representative electropherogram and pseudo-gel to study the effect of **1** in WT myoblasts. (**C**) Representative electropherogram and pseudo-gel to study the effect of **1** in DM1 patient-derived myoblasts. In panels B and C, exon 25 inclusion is indicated by "+E" while exclusion is indicated by "-E" (n = 3 biological replicates). The additional band above "+E" represents a hybrid duplex formed by annealing between the exon5–containing and exon5–skipped amplicons and was excluded from quantification.

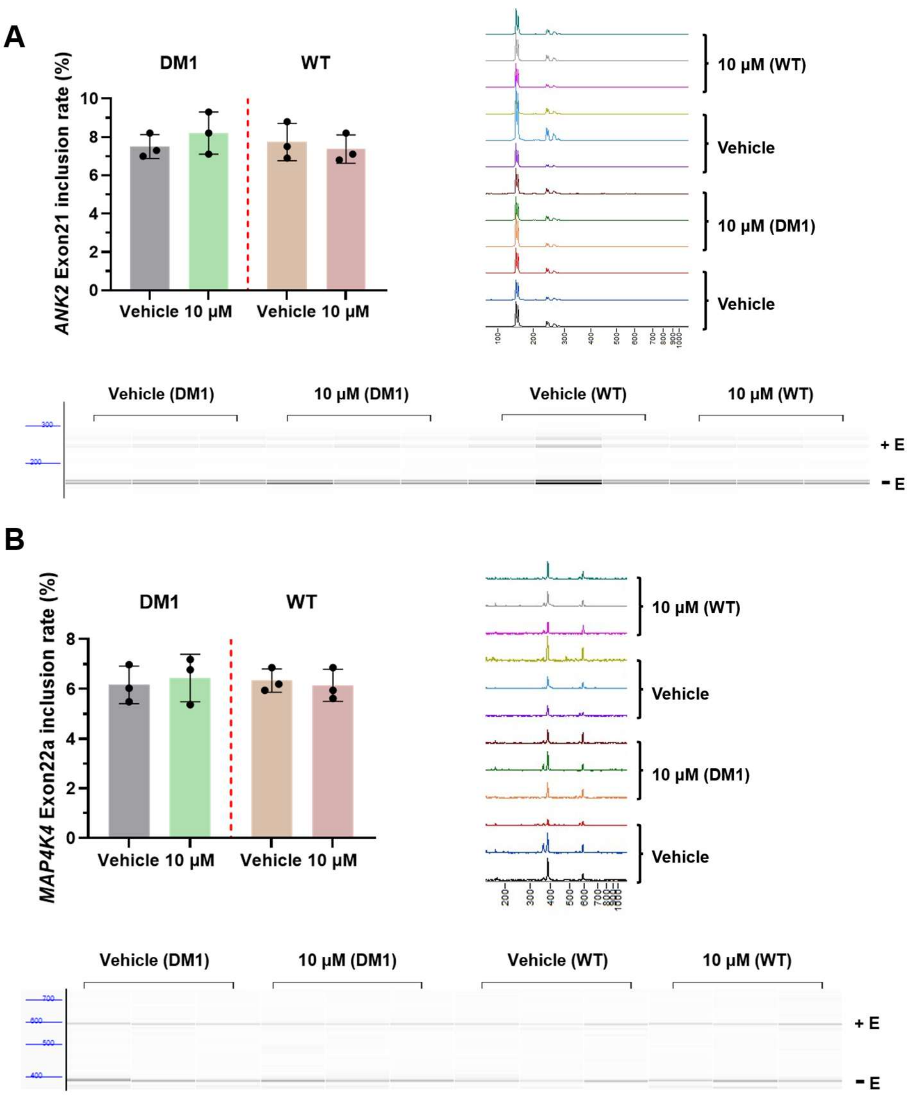

**Fig. S6. Effect of 1 on MBNL1-independent splicing events in DM1 and WT myoblasts.** (A) Effect of **1** (10 μM) on NOVA-regulated *MAP4K4* exon 22a splicing, assessed by end-point RT-PCR and fragment analyzer (n = 3 biological replicates), including a representative electropherogram and pseudo-gel. Exon 22a inclusion is indicated by "+E" while exclusion is indicated by "-E". (B) Effect of **1** (10 μM) on *ANK2* exon 21 splicing events assessed by end-point RT-PCR and fragment analyzer (n = 3 biological replicates), including a representative electropherogram and pseudo-gel. Exon 21 inclusion is indicated by "+E" while exclusion is indicated by "-E".

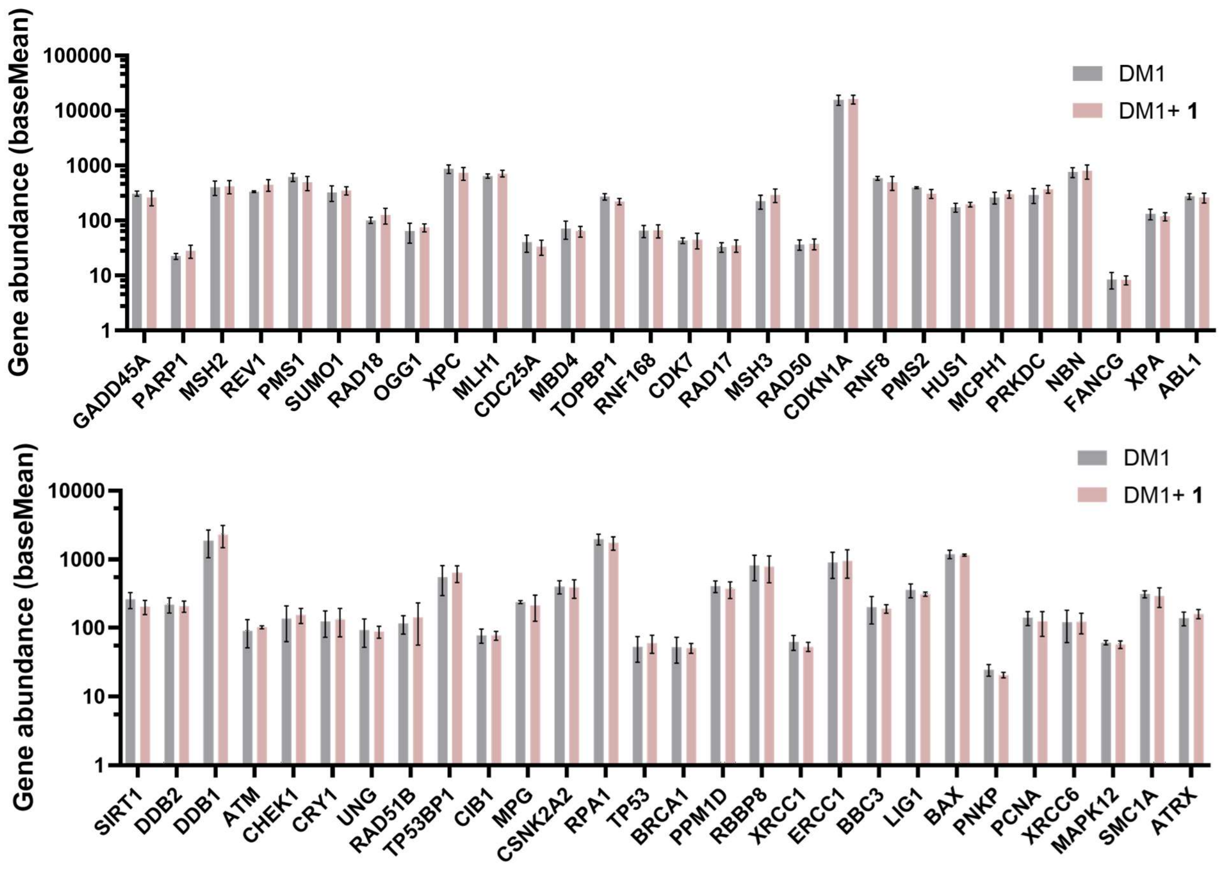

**Fig. S7. RNA sequencing analysis of potential off-targets in the DNA damage pathway in DM1 myoblasts treated with 1.** RNA-seq analysis of 56 genes involved in DNA damage response pathways [6] in DM1 myoblast treated with 10 µM of **1** or 0.1% (v/v) DMSO (n = 2 biological replicates). Data are reported as the mean ± SD.

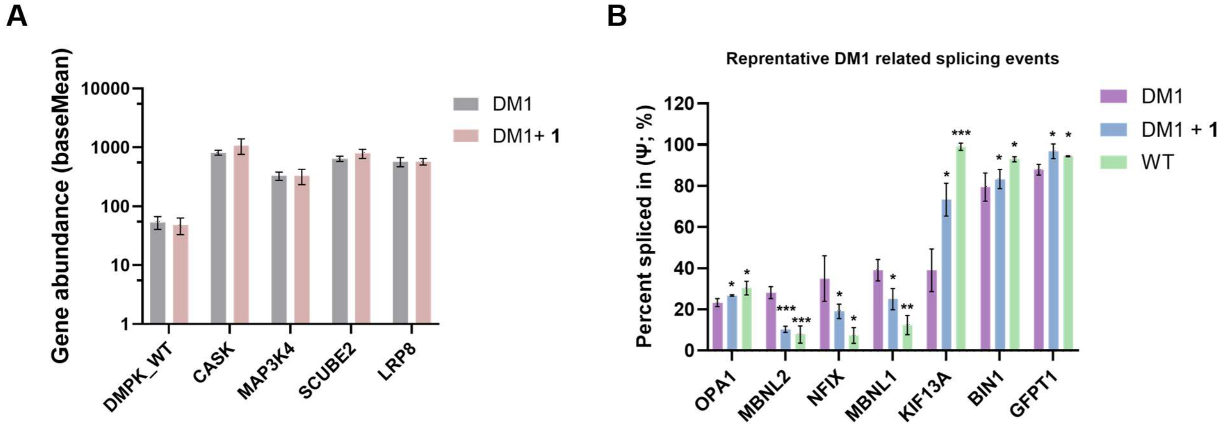

**Fig. S8. Transcriptome-wide analysis of transcripts with short r(CUG) repeats and DM1-associated splicing event upon treatment of DM1 myoblasts with 1 (10 μM).** (**A**) Analysis of gene abundance of transcripts containing short, non-pathogenic r(CUG) repeats in DM1 myoblasts treated with 10 μM **1** or 0.1% (v/v) DMSO from RNA-seq studies (n = 2 biological replicates). Data are reported as the mean ± SD. (**B**) Analysis of seven representative splicing events previously reported to have correlation with DM1 molecular pathology and clinical manifestations [7-8]. Data are reported as the mean ± SD. (**) $P < 0.01$, (***) $P < 0.001$, as determined by comparison to vehicle-treated DM1 myoblasts (0.1% (v/v) DMSO) by SUPPA2 [9].

#### EXPERIMENTAL METHODS

##### Biochemical Methods

***In vitro* transcription of r(CUG)$_{42}$, r(CUG)$_{20}$ and r(CAG)$_{20}$:** The DNA template for *in vitro* transcription of r(CUG)$_{42}$ repeats was obtained by amplifying the desired sequence from a kanamycin-resistant pCTG109 vector, which was previously cloned from a pCNA plasmid containing the repeat expansion sequence [10]. The r(CUG)$_{20}$ and r(CAG)$_{20}$ were obtained from the template in Table **S1**. Optimal PCR conditions were achieved using 35 cycles (denaturing at 95 ºC for 30 s, annealing at 61 °C for 30 s, extension at 72 ºC for 45 s) and a final elongation at 72 ºC for 7 min, using 50 nM of DNA template, 1 µM each of forward and reverse primers, 1 mM of each dNTP, 8% (v/v) DMSO, 1× Phusion HF buffer, and a Phusion™ High-Fidelity DNA Polymerase concentration of 0.4 U/µL (New England Biolabs, MA, USA). After amplification, the plasmid DNA was digested using DpnI, and the enzyme was heat-inactivated by incubating at 80 ºC for 20 min, according to the manufacturer's instructions (New England Biolabs, MA, USA).

Transcription of r(CUG)$_{42}$ was completed in a 4 mL reaction containing 40 mM Tris, pH 8, 1 mM spermidine, 0.01% (v/v) Triton X-100, 5 mM dithiothreitol (DTT), 3 µL of PCR product, 24 mM $MgCl_2$, 16 mM rNTPs mixture, 8% (v/v) DMSO, and 0.07 mg/mL of in-house prepared and purified T7 polymerase. The reaction was incubated for 4 h at 37 °C.

The fully base paired RNA control was formed by annealing r(CUG)$_{20}$ and r(CAG)$_{20}$. Each RNA was *in vitro* transcribed from a ssDNA template obtained from Integrated DNA Technologies (IDT. For each RNA, *in vitro* transcription was completed in a solution containing 40 mM Tris, pH 8, 1 mM spermidine, 0.01% (v/v) Triton X-100, 5 mM DTT, 1.6 µM of T7 promoter and ssDNA, 55 mM $MgCl_2$, 32 mM rNTPs mixture, 8% (v/v) DMSO, 8% (v/v) PEG, and 0.07 mg/mL of in-house prepared and purified T7 polymerase. The reaction was incubated for 4 h at 37 °C.

The RNAs were purified by gel electrophoresis using denaturing 10% (w/v) (r(CUG)$_{42}$) or 15% (w/v) 19:1 for the fully base paired control RNA (29:1 acrylamide:bisacrylamide) gels. The desired band was visualized by UV shadowing and extracted from the gel by passive elution of the crushed gel in 0.3 M sodium acetate, pH 5, at 4 °C. The supernatant was then ethanol precipitated twice and desalted using Sephadex G-25 cartridges (Cytiva) per the manufacturer's recommended protocol.

***In vitro* cleavage assays.** The $r(CUG)_{12}$ RNA (1 µM; purchased from Dharmacon) was folded in 1× Assay Buffer (20 mM HEPES, 25 mM KCl, pH 7.5) by heating at 90°C for 3 min followed by snap cooling on ice for 10 min. The $r(CUG)_{42}$ and fully paired RNA control were folded at 0.25 µM in 1× Assay Buffer following the same protocol. The concentration adjustment allows for targeting a similar number of repeats within two RNAs different in length, where $r(CUG)_{12}$ has five 1×1 UU internal loops and $r(CUG)_{42}$ has 20. Compounds were pre-complexed with freshly made 1.0 mol eq. $Fe(II)(NH_4)_2(SO_4)_2$ solution (typically a 5 mM aqueous solution of $Fe(II)(NH_4)_2(SO_4)_2$ was mixed with 5 mM DMSO solution of the compound) and were further diluted to the assay concentrations in 1× Assay Buffer. Reactions were assembled in a 384-well transparent plate (Fisher Scientific, catalog #12565506, typically 100 µL reaction final volume), and lastly a freshly made 10 mM solution of DTT was added to a final concentration of 1 mM. (When hydrogen peroxide was used in the reaction, a master mix of 1 mM DTT and 1 mM $H_2O_2$ in water was prepared and added.) The plate was incubated at 37°C for 30 min, 2 h and 6 h with gentle shaking. The reaction mixtures were separated by gel electrophoresis using a denaturing 8% or 20% (w/v) polyacrylamide gel (RNA Ladders were purchased from New England Biolabs, catalog #N0364S and #N2102S). The gel was then post-stained with SYBR Gold solution (1:10,000 dilution in 1× TBE buffer) for 5 min and imaged using an Azure Biosystems Sapphire Biomolecular or Amersham Typhoon (Cytiva) imager. The bands were quantified using Image J 3, and graphs were generated using Prism 10.

#### Biology Methods

**Cell Lines.** DM1 conditional MyoD-fibroblast cells and wild-type conditional MyoD-fibroblast cells [11] (gifts from D. Furling; Centre de Recherche en Myologie (UPMC/Inserm/CNRS), Institut de Myologie, Paris France) were maintained in Dulbecco's Modified Eagle Medium (DMEM; Corning, catalog #15-017-CV) supplemented with 15% (v/v) fetal bovine serum (FBS; Gibco, catalog #10437-028), 1% (v/v) Antibiotic-Antimycotic Solution (Corning; #30-004-CI), and 1% (v/v) Glutagro (Corning, catalog #25-015-CI) in 100 mm diameter dishes.

For compound treatment, cells were seeded into 24-well plates and cultured in growth medium until ~100% confluent. Conditional MyoD-fibroblast cells were then differentiated into myoblasts using a differentiation medium composed of 1× DMEM, 1% (v/v) Antibiotic-Antimycotic Solution, 0.1 mg/mL transferrin human (Sigma, #T8158), 0.01 mg/mL insulin (Sigma, catalog #I0516), and

2 μg/mL doxycycline (freshly added; Sigma, catalog #10592-13-9). After the initial 24 h differentiation period, cells were treated with the compound of interest (0.1% (v/v) DMSO) for an additional 24 h in the same differentiation medium. Cells were used with passage numbers less than 20.

HeLa480 cells [1] were maintained in DMEM (Corning, catalog #15-017-CV) supplemented with 10% (v/v) FBS (Gibco, catalog #12676029), 1% (v/v) Antibiotic-Antimycotic Solution (Corning, catalog #30-004-CI), and 1% (v/v) Glutagro (Corning, catalog #25-015-CI). These cells were cultured at 37 °C in a humidified atmosphere containing 5% $CO_2$ with passage numbers less than 20.

**RT-qPCR Analysis.** DM1 conditional MyoD-fibroblasts and HeLa480 were cultured and treated in 24-well plates as described above. Total RNA was harvested using a Quick-RNA Mini Prep Kit (Zymo Research) following the manufacturer's recommended protocol, including the on-column DNase I digestion. Approximately 300 ng of total RNA was reverse transcribed in a 20 μL reaction using a qScript cDNA Synthesis Kit (Quanta BioSciences) per the manufacturer's recommended protocol. Then, qPCR analysis was carried out with 2 μL of the RT reaction and the corresponding pair of primers (Table S1) and SYBR Green Master Mix (Thermo Fisher Scientific, catalog #A25741) on a QuantStudio 5, 384-well Block Real-Time PCR System (Applied Biosystems). Amplification conditions were 95°C for 2 min, followed by 40 cycles of 95°C for 15 s and 60°C for 1 min. Relative abundance of each transcript was determined by normalizing to the housekeeping gene (*GAPDH*) using the $2^{-\Delta\Delta Ct}$ method [12].

**Evaluation of Alternative pre-mRNA Splicing by End-Point RT-PCR Analysis [13].** End-point PCR reactions were carried out with cDNA samples from "**RT-qPCR Analysis**" by using GoTaq Flexi DNA Polymerase kit (Promega) per the vendor's recommendations and following cycling conditions: 30 cycles of 95 °C for 30 s, 58 °C for 30 s, and 72 °C for 1 min and a final extension at 72 °C for 5 min using primers listed in Table S1. The amplified PCR products were analyzed and quantified using dsDNA 910 Reagent Kit (35–1500 bp, catalog #DNF910K0500) by 5300 Fragment Analyzer System (Agilent Technologies). Fluorescence data generated during capillary electrophoresis were processed by instrument software to produce electropherograms and pseudo-gel. DNA concentrations were calculated automatically from integrated peak areas across and calibrated according to fragment size by Fragment Analyzer software. The resulting concentration measurements were used to determine the relative proportions of exon-included (+E) and exon-skipped (−E) splice products.

Exon inclusion rates were calculated using the following equation:

$$\%exon\ inclusion\ rate = 100 \times \frac{Signal\ from\ band\ where\ exon\ is\ included}{Signal\ from\ band\ where\ exon\ is\ included + Signal\ from\ band\ where\ exon\ is\ excluded}$$

(Equation 1)

Percent rescue was calculated using the following equation:

$$\%\ Rescue = 100 \times \frac{\%\ exon\ inclusion\ untreated\ DM1\ cells - \%\ exon\ incusion\ treated\ DM1\ cells}{\%\ exon\ inclusion\ untreated\ DM1\ cells - \%\ exon\ inclusion\ untreated\ WT\ cells}$$

(Equation 2)

**Cellular NanoBRET Assay to study r(CUG)$^{exp}$-MBNL1 complex formation[4]**. HeLa480 cells were seeded into 6-well plates and co-transfected with 2.5 μg MBNL1-HaloTag and 50 ng MBNL1-NanoLuc plasmids using 8 μL of FuGENE® HD Transfection Reagent (Promega, catalog #E2311) in 110 μL of Opti-MEM (Gibco, catalog #31985070), following the manufacturer's instructions. After 16 h, cells were resuspended in an appropriate volume of assay medium (Opti-MEM + 4% (v/v) FBS) and re-seeded into white 96-well tissue culture plates (Corning, catalog #3903) with $1.6 \times 10^4$ transfected cells per well (affording ~70% confluency). Each well received 100 μL of growth medium with 200 nM HaloTag 618 Ligand (Promega, catalog #N1662), while three control wells without ligand were prepared. After incubating for 4 h at 37 °C, the compound of interest (or vehicle) was added at the desired concentration with a final DMSO concentration of 0.1% (v/v). Three wells were treated with 10 μM CAG25 vivo morpholino as positive control.

The following day, NanoLuc substrate and extracellular NanoLuc inhibitor were added. For a 96-well plate, 2.5 mL of Opti-MEM were combined with 25 μL of NanoLuc substrate (Promega, catalog #N1662) and 8.3 μL of extracellular NanoLuc inhibitor (Promega, catalog #N2160) to afford a 5× working solution. Then, 25 μL of working solution was added to each well. BRET measurements were then collected on a GloMax Discover System (Promega, catalog #GM3000). Dual-filtered luminescence was collected with a 460/80 nm bandpass filter for the NanoLuc donor and a 610 nm long-pass filter for the HaloTag acceptor using an integration time of 500 ms. The corrected NanoBRET ratio, was calculated using Equation 3 with milliBRET units (mBU):

$$\text{Corrected NanoBRET Ratio} = \frac{\text{HaloTag ligand (Emission 610 nm/LP)}}{\text{Donor Luminescence (Emission 450 nm/BP 80 nm)}} - \frac{\text{No Ligand Control (Emission 610 nm/LP)}}{\text{Donor Luminescence (Emission 450 nm/BP 80 nm)}}$$

(Equation 3)

**Evaluation of γ-H2AX Expression via TR-FRET**. HeLa480 cells were plated in a 96-well plate and treated with compound of interest in growth medium for 24 h. After 24 h, the FRET signal emanating from γ-H2AX foci were measured using a HTRF Human Phospho-H2AX (Ser139) Detection Kit (Revvity, catalog #64H2XPEG) according to the manufacturer's protocol.

**Evaluation of γ-H2AX Expression via Immunostaining**. DM1 fibroblasts were differentiated and treated in 24-well plates containing 12 mm glass cover slips (Fisher Scientific, catalog # NC1418755) with the compounds of interest as described in "**Cell Lines**". Cells were fixed with 4% (w/v) paraformaldehyde for 10 min at 37 °C, followed by overnight permeabilization in 75% (v/v) ethanol at 4 °C. After permeabilization, cells were washed with 1× DPBS and incubated overnight at 4 °C with Alexa Fluor 488 Anti-gamma H2A.X (phospho S139) antibody (Abcam, catalog #ab195188) diluted 1:1000 in 1× PBS. After incubation, samples were mounted in 20 µL ProLong Gold Antifade Mountant with DAPI (Invitrogen, catalog #P36931) and imaged on a Zeiss LSM 980 Airyscan 2 confocal microscope with oil immersion at 63x magnification.

**Evaluation of r(CUG)$^{exp}$ Foci Containing MBNL1 via smFISH and Immunostaining**. Single molecule fluorescence *in situ* hybridization (smFISH) probes targeting *DMPK* coding sequence (CDS) were used as previously designed with sequences provided in Table S2 [4]. Steps for compound treatment, fixation, and permeabilization were performed as described in "**Evaluation of γ-H2AX Expression via Immunostaining**". After permeabilization, cells were washed with 1× DPBS and processed for imaging MBNL1 (immunofluorescence) and then r(CUG)$^{exp}$ (smFISH). For endogenous MBNL1 imaging, cells were incubated overnight at 4 °C with Anti-MBNL1 Monoclonal Antibody (EMD Millipore, catalog #MABE70) diluted 1:1000 in 1× PBS. After washing with 1x DPBS three times, samples were incubated with Alexa Fluor 488 goat anti-mouse IgG (H+L) (invitrogen, catalog # A-11001, 1:2000 dilution in 1× PBS) for 2 h at room temperature in the dark.

Next, smFISH *DMPK* probes (100 nM each) conjugated to ddUTP-Atto550 (Axxora, catalog #JBS-NU-1619-550) were prepared in 1× Hybridization Buffer (2× SSC, 10% (w/v) dextran sulfate, 10% (v/v) formamide) and added to cells in a humidified chamber overnight at 37 °C. Cells were washed three times with 2× SSC at room temperature to remove excess probes. Finally, samples were mounted in 20 μL ProLong Gold Antifade Mountant with DAPI (Invitrogen, catalog #P36931) and imaged on a Zeiss LSM 980 Airyscan 2 confocal microscope with oil immersion at 63x magnification.

**RNA-Seq**. The quality of RNA extracted from "**RT-qPCR Analysis"** were checked by an Agilent 2100 Bioanalyzer Nano chip and quantified using a Qubit 2.0 Fluorometer (Invitrogen). Sequencing libraries were prepared using the cDNA-PCR Sequencing Kit V14 with barcoding (Oxford Nanopore Technologies, catalog #SQK-PCB114.24) following the manufacturer's instructions. The final libraries were loaded onto a PromethION 48 instrument equipped with R10.4.1 flow cells (Oxford Nanopore Technologies, catalog #FLO-PRO114M). Nanopore raw signals (POD5) were base called using MinKNOW software on PromethION with the high-accuracy model and aligned to a human reference genome (GRCh38) by minmap2 [14]. For transcriptome-wide analysis, gene-level differences were quantified using DESeq2 [15], based on normalized read counts generated by StringTie2 [16]. Genes with baseMean < 10 were excluded, and genes with $\log_2$ fold change > 0.5 and FDR < 0.05 were considered differentially expressed. PSI quantification and differential splicing analysis were performed using SUPPA2 (v2.4, https://github.com/comprna/SUPPA) [9]. Splicing events with |ΔPSI| > 0.1 and FDR < 0.05 were classified as significantly altered.

#### SYNTHETIC METHODS

**General Synthetic Methods**. Chemicals were purchased from Sigma Aldrich, ChemScene, CombiBlocks, or BroadPharm and were used without further purification. All reactions involving air- or moisture-sensitive reagent were performed under argon atmosphere.

Thin layer chromatography (TLC) on silica gel was carried out on Sigma-Aldrich pre-coated silica plates with 0.25 mm layer thickness to monitor the progress of the reactions. Compounds were visualized by irradiation (254 nm) and/or staining with phosphomolybdic acid, vanillin or ninhydrin stains. Anhydrous $MgSO_4$ was used as a drying agent for organic phases. Reaction progress was also monitored by liquid chromatography-mass spectrometry (LC-MS) on Agilent 1260 Infinity II system equipped with 1260 Infinity II binary pump, 1260 Infinity II diode array detector (DAD), 1260 Infinity II Multisampler and 6130 Single Quadruple LC-MS (electrospray ionization; ESI). Agilent SB-C18 1.8 μm OBD, 2.1 ×50 mm column was used with a linear gradient of 5-95% methanol (MeOH) in water with 0.1% (v/v) formic acid (FA) and a flow rate of 0.350 mL/min, applied over 11 min.

Flash chromatography was performed on silica gel columns (Silicycle® UltraPure SiliaSep™ 230-400 mesh 40-63 μM, 4-40 g) using a Biotage® Isolera One purification system with a variable flow depending on the column size. Reverse phase preparative high performance liquid chromatography (HPLC) was performed using a Waters system equipped with 1525 Binary HPLC pump, a Waters 2487 dual absorbance detector system and a Waters Sunfire C18 OBD 5 μm 19 x 150 mm column. Absorbance was monitored at 254 and 220 nm. A gradient of 0-100% MeOH in water with 0.1% (v/v) trifluoroacetic acid (TFA) over 60 min with a flow rate of 5 mL/min was used for purification of synthesized compounds. Purity was assessed by analytical HPLC using a Waters Symmetry C18 5 μm, 4.6 × 150 mm column with a flow rate of 1 mL/min and a linear gradient from 0-100% MeOH in water with 0.1% (v/v) TFA applied over 70 min; absorbance was monitored at 254 nm. All compounds used for biochemical or biological evaluation are >95% pure by HPLC analysis.

1H NMR and 13C NMR spectra were collected on a 400 MHz UltraShieldTM equipped with cryo probe. Chemical shifts ($\delta$) are reported in parts per million (ppm) referenced to the residual 1H resonance of the solvent (DMSO $\delta$ 2.50). The following abbreviations are used for coupling patterns: br=broad, s=singlet, d=doublet, dd=doublet of doublets, t=triplet, q=quartet, quint=quintuplet, m=multiplet, dt, doublet of triplets, dq=doublet of quartets, td=triplet of doublets. Coupling constants (J) are listed in hertz (Hz). Due to the interconversion of conformers at room temperature, the 1H NMR spectra of compound **3** was collected at 80 °C.

Low resolution MS spectra were obtained with a LC-MS on Agilent 1260 Infinity II system equipped with 1260 Infinity II binary pump, 1260 Infinity II DAD, 1260 Infinity II Multisampler and 6130 Single Quadruple LC-MS (ESI). High resolution mass spectrometry was carried out using a LTQ Orbitrap hybrid mass spectrometer with an electrospray ionization probe (Thermo Scientific, San Jose, CA) by direct infusion from the pump syringe.

Intermediates 4-(3-(6-(4-methylpiperazin-1-yl)-1H,3'H-[2,5'-bibenzo[d]imidazol]-2'-yl)phenoxy)-butanoic acid and compound **3** were synthesized following previously reported protocols [17-19].

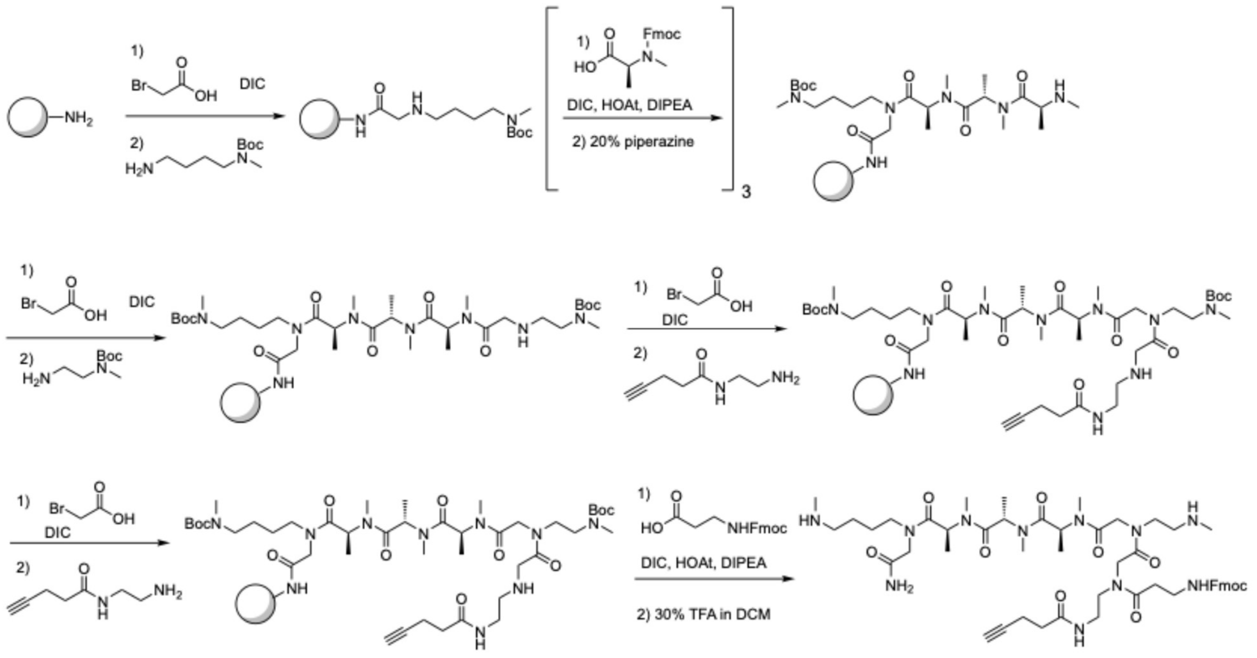

**Scheme 1. Solid phase synthesis of linker intermediate 4.**

**Compound 4**. Compound **4** was synthesized by adapting a previously reported protocol (**Scheme 1**) [20]. For all microwave reactions, a 700 W microwave set to 10% power was used. Rink amide resin (1 g, 0.6 mmol) was swollen in *N*,*N*-dimethyl formamide (DMF) at room temperature for 10 min and then deprotected with a solution of 20% piperidine in DMF (5 mL, 2 × 20 min). The resin was washed with DMF (3 × 5 mL) and reacted twice with a solution of 2 M bromoacetic acid (4 mL) in DMF and *N,N'*-diisopropylcarbodiimide (DIC; 500 µL, 3.2 mmol) via microwave irradiation

(3 × 15 s). The resin was washed with DMF (3 × 5 mL) and reacted with a solution of *N*-(4-aminobutyl)-*N*-methyl carbamic acid tert-butyl ester (240 mg, 1.2 mmol) in DMF (4 mL) via microwave irradiation (3 × 15 s).

The resin was washed with DMF and reacted twice with a solution of 2 M bromoacetic acid (4 mL) in DMF and DIC (500 μL, 3.2 mmol) via microwave irradiation (3 × 15 s). The resin was washed with DMF (3 × 5 mL) and reacted with a solution of tert-butyl (2-aminoethyl)(methyl)carbamate) (210 mg, 1.2 mmol) in DMF (4 mL) via microwave irradiation (3 × 15 s). The resin was shaken with this solution at room temperature for an additional 1 h. Next, the resin was washed with DMF (3 × 5 mL) and reacted twice with a solution of 2 M bromoacetic acid (4 mL) in DMF and DIC (500 μL, 3.2 mmol) via microwave irradiation (3 × 15 s).

The resin was washed with DMF (3 × 5 mL) and reacted with a solution of *N*-(2-aminoethyl)pent-4-ynamide (170 mg, 1.2 mmol) in DMF (4 mL) via microwave irradiation (3 × 15 s). The resin was shaken with this solution at room temperature for an additional 1 h, washed with DMF (3 × 5 mL) and reacted with a solution of Fmoc-β-alanine (500 mg, 1.62 mmol), DIC (500 μL, 3.2 mmol), 1-hydroxy-7-azabenzotriazole (HOAt; 220 mg, 1.62 mmol), and *N*,*N*-diisopropylethylamine (DIEA; 280 μL, 1.62 mmol) in DMF (4 mL) via microwave set to 75 °C for 10 min.

Lastly the resin was washed with DMF and DCM and then treated with 30% (v/v) TFA/DCM (10 mL) for 10 min. The solution was concentrated *in vacuo* and co-evaporated with toluene three times to afford a pale yellow oil of **4** that was used in the next step without further purification. **MS (ESI)** m/z = 502.0 $[M+2H]^{2+}$.

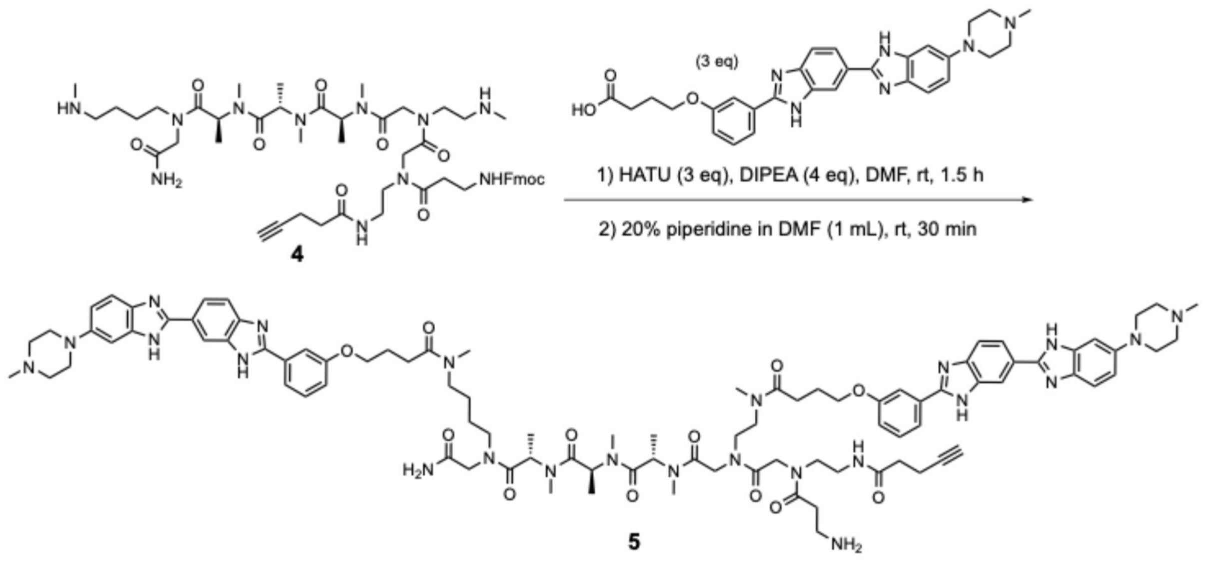

**Scheme 2. Synthesis of 5.**

**Compound 5**. To a stirred solution of 4-(3-(6-(4-methylpiperazin-1-yl)-1H,3'H-[2,5'-bibenzo[d]imidazol]-2'-yl)phenoxy)butanoic acid (60 mg, 0.12 mmol, 3 eq, synthesized as previously reported) in DMF (3 mL), hexafluorophosphate azabenzotriazole tetramethyl uronium (HATU; 45 mg, 0.12 mmol, 3 eq) and DIEA (28 μL, 0.16 mmol, 4 eq) were added [17]. The reaction mixture was stirred for 15 min at room temperature. Next, **4** (40 mg, 0.04 mmol, 1 eq) was added and the reaction mixture was stirred for 1.5 h at room temperature. The progress of the reaction was monitored by LC-MS. After completion, the crude mixture was concentrated under reduced pressure and purified by reverse phase prep-HPLC as described in the **General Synthetic Methods**. Fmoc was removed by treating with 1 mL of 20% (v/v) piperidine/DMF (1 x 10 min) and then purified by reverse phase prep-HPLC as described in the **General Synthetic Methods** to afford compound **5** as a yellow solid. Yield: 32%, 22 mg. **MS (ESI)** m/z = 442.3 $[M+4H]^{4+}$.

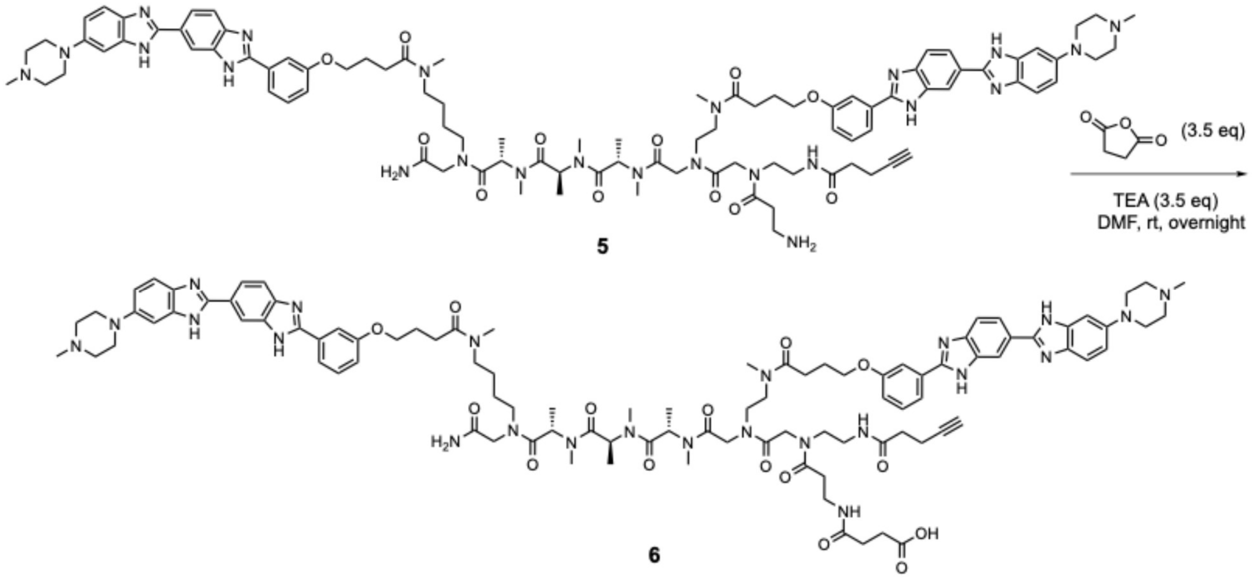

**Scheme 3. Synthesis of 6.**

**Compound 6**. To a stirred solution of **5** (10 mg, 5.7 μM, 1 eq) in DMF, succinic anhydride (2 mg, 0.02 mmol, 3.5 eq) and triethylamine (TEA; 3 μL, 0.02 mmol, 3.5 eq) were added. The reaction mixture was stirred at room temperature overnight. The progress of the reaction was monitored by LC-MS. After completion, the crude mixture was concentrated under reduced pressure and purified by reverse phase prep-HPLC as described in the **General Synthetic Methods** to afford compound **6** as a yellow solid. Yield: 80%, 8.5 mg. **MS (ESI)** m/z = 622.3 $[M+3H]^{3+}$.

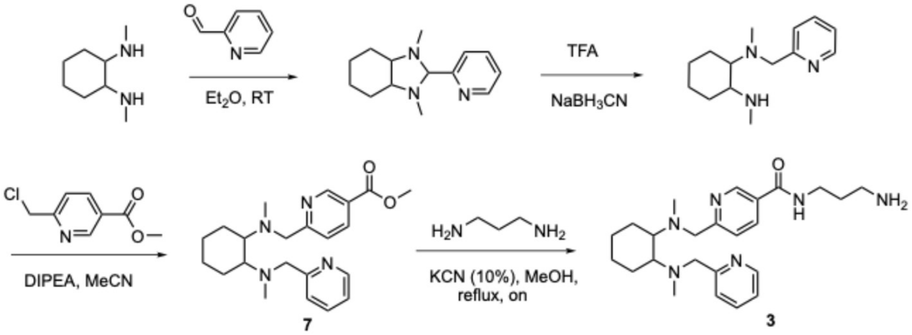

**Scheme 4. Synthesis of tetradentate pyridine Fe(II) chelating module 3.**

**Methyl6-((methyl(2-(methyl(pyridin-2-yl-methyl)amino)cyclohexyl)amino)methyl)nicotinate (compound 7).** To a stirring solution of *N*[1],*N*[2]-dimethyl-N[1]-(pyridin-2-ylmethyl)cyclohexane-1,2-diamine (89 mg, 0.28 mmol, 1.0 eq, synthesized as previously described [18]) in acetonitrile (MeCN) under argon atmosphere, methyl 6-(chloromethyl)nicotinate (80 mg, 0.28 mmol, 1.0 eq, synthesized as previously described [19]) and DIEA (100 µL) were added. The solution was stirred under reflux overnight, and conversion was monitored via LC-MS. When compete conversion of the starting material was observed, the solvent was evaporated under reduced pressure and the crude material was used in the next step without further purification. **MS (ESI)** m/z = 383.2 $[M+H]^+$.

***N*-(3-aminopropyl)-6-((methyl(2-(methyl(pyridin-2-ylmethyl)amino)cyclohexyl)amino)-methyl)nicotinamide (compound 3).** Compound **3** was synthesized by adapting a previously reported protocol [19]. To a stirring solution of **7** in MeOH (76 mg, 0.20 mmol, 1.0 eq), propane-1,3-diamine (170 µL, 2.0 mmol, 10 eq) and KCN (10% mol) were added. The resulting mixture was stirred at reflux for 24 h and conversion was monitored by LC-MS. After completion, the crude mixture was concentrated under reduced pressure and purified by reverse phase prep-HPLC as described in the **General Synthetic Methods** to afford compound **3** as a white solid. Yield: 35%, 30 mg; **$^1$H NMR** (DMSO-$d_6$, 400 MHz) δ: 8.89 (d, *J* = 4.0 Hz, 1H), 8.62 (t, *J* = 4.0 Hz, 1H), 8.49 (q, *J* = 4.0 Hz, 1H), 8.23 (dd, *J* = 9.1, 2.3 Hz, 1H), 7.98 (td, *J* = 7.7, 1.8 Hz, 1H), 7.58 (d, *J* = 8.1 Hz, 1H). 7.49 (d, *J* = 7.8 Hz, 1H), 7.40-7.37 (m, 1H), 4.39 (d, *J* = 8.0 Hz, 1H), 4.7 (d, *J* = 8.0 Hz, 1H), 4.17-3.91 (m, 2H), 3.39 (q, *J* = 4.0 Hz, 2H), 3.35-3.29 (m, 1H), 3.28-3.17 (m, 1H), 2.90 (s, 2H), 2.83 (s, 3H), 2.48 (s, 2H), 2.15 (t, *J* = 8.0 Hz, 2H), 1.94-1.75 (m, 4H), 1.61-1.41 (m, 2H), 1.41-1.19 (m, 2H). **MS (ESI)** m/z = 425.3 $[M+H]^+$. **HR-MS (ESI)**, m/z calculated for $C_{24}H_{37}N_6O$ [M+H]+ 425.3034, found 425.3021. **$HPLC_{RT}$** : 11.6 min. **HPLC purity**: 95 %.

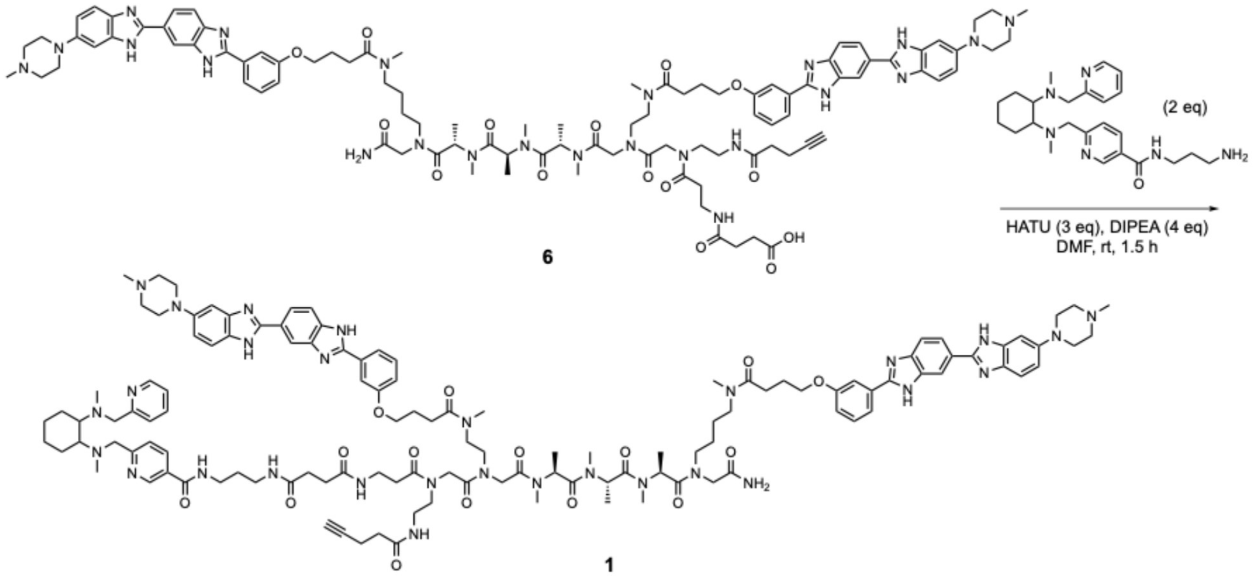

**Scheme 5. Synthesis of heterobifunctional 1.**

**Compound 1.** To a stirred solution of **6** (4 mg, 2.14 μM, 1 eq) in DMF (1 mL), HATU (2.45 mg, 6.43 μM, 3 eq) and DIEA (1.5 μL, 8.56 μM, 4 eq) were added. The reaction mixture was stirred for 15 min at room temperature. Next, **3** (1.82 mg, 4.28 μM, 2 eq) was added, and the reaction mixture was stirred for 1.5 h at room temperature. The progress of the reaction was monitored by LC-MS. After completion, the crude mixture was concentrated under reduced pressure and purified by reverse phase prep-HPLC as described in the **General Synthetic Methods** to afford target compound **1** as a light-yellow solid. Yield: 24%, 1.17 mg. **HR-MS (ESI)**, *m/z* calculated for $C_{122}H_{159}N_{29}O_{15}$ $[M+5H]^{5+}$ 455.2535, found 455.2509. **HPLC$_{RT}$** : 32.4 min. **HPLC purity**: 95.6 %.

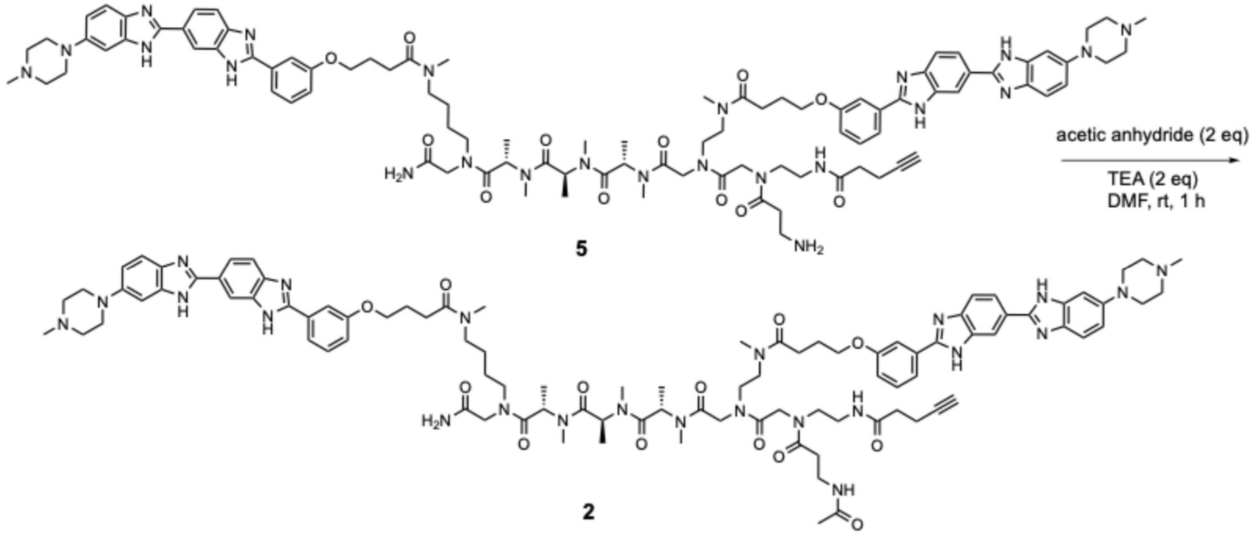

**Scheme 6. Synthesis of binder 2.**

**Compound 2**. To a stirred solution of **5** (5 mg, 30 μM 1 eq) in DMF (1 mL), acetic anhydride (0.6 μL, 60 μM, 2 eq) and TEA (0.8 μL, 60 μM, 2 eq) were added. The reaction mixture was stirred at room temperature for 1 h. The progress of the reaction was monitored by LC-MS. After completion, the crude mixture was concentrated under reduced pressure and purified by reverse phase prep-HPLC as described in the **General Synthetic Methods** to afford **2** as a light-yellow solid. Yield: 36 %, 1.9 mg. **HR-MS (ESI)**, *m/z* calculated for $C_{96}H_{123}N_{23}O_{13}$ $[M+4H]^{4+}$ 452.4923, found 452.7489. **HPLC$_{RT}$** : 29.0 min. **HPLC purity**: 96.7 %.

### COMPOUND CHARACTERIZATION

#### Compound 1:

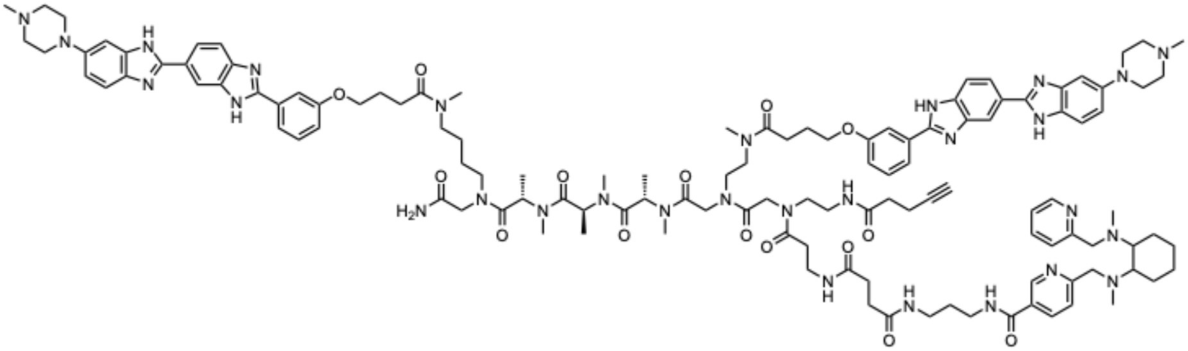

## HR-MS:

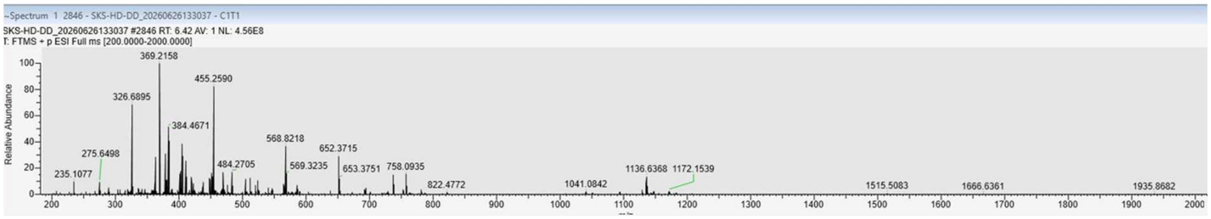

#### HPLC:

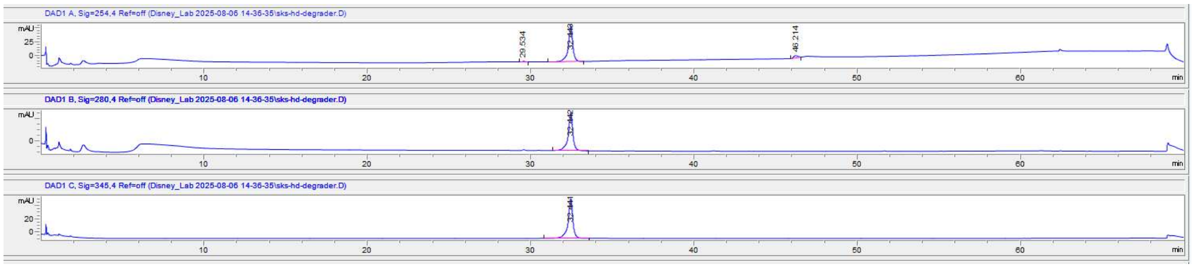

#### Compound 2:

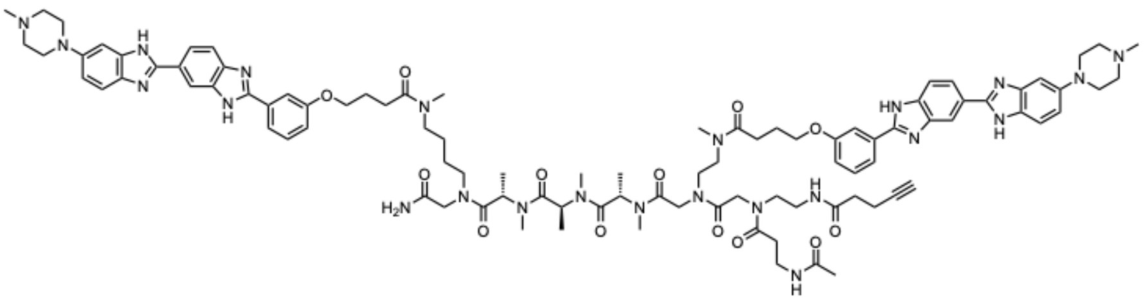

## HR-MS

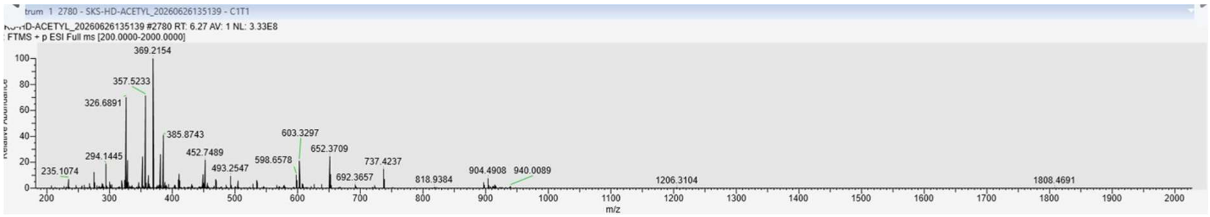

#### HPLC:

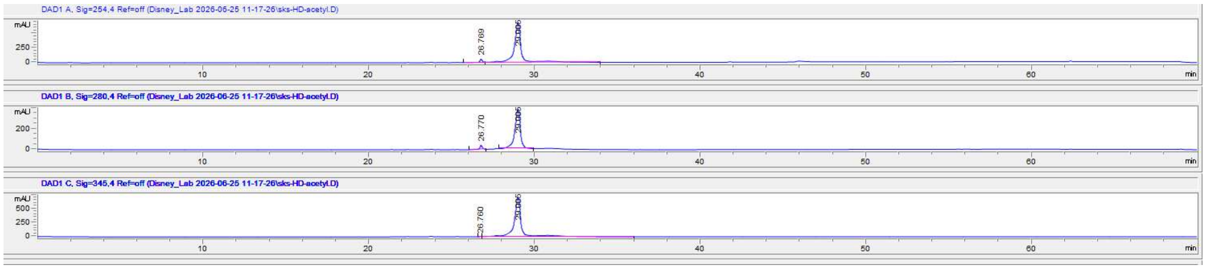

**N-(3-aminopropyl)-6-((methyl(2-(methyl(pyridin-2-ylmethyl)amino)cyclohexyl)amino)-methyl)nicotinamide (3):**

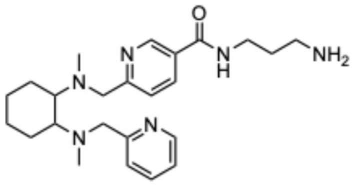

**$^{1}$H NMR**

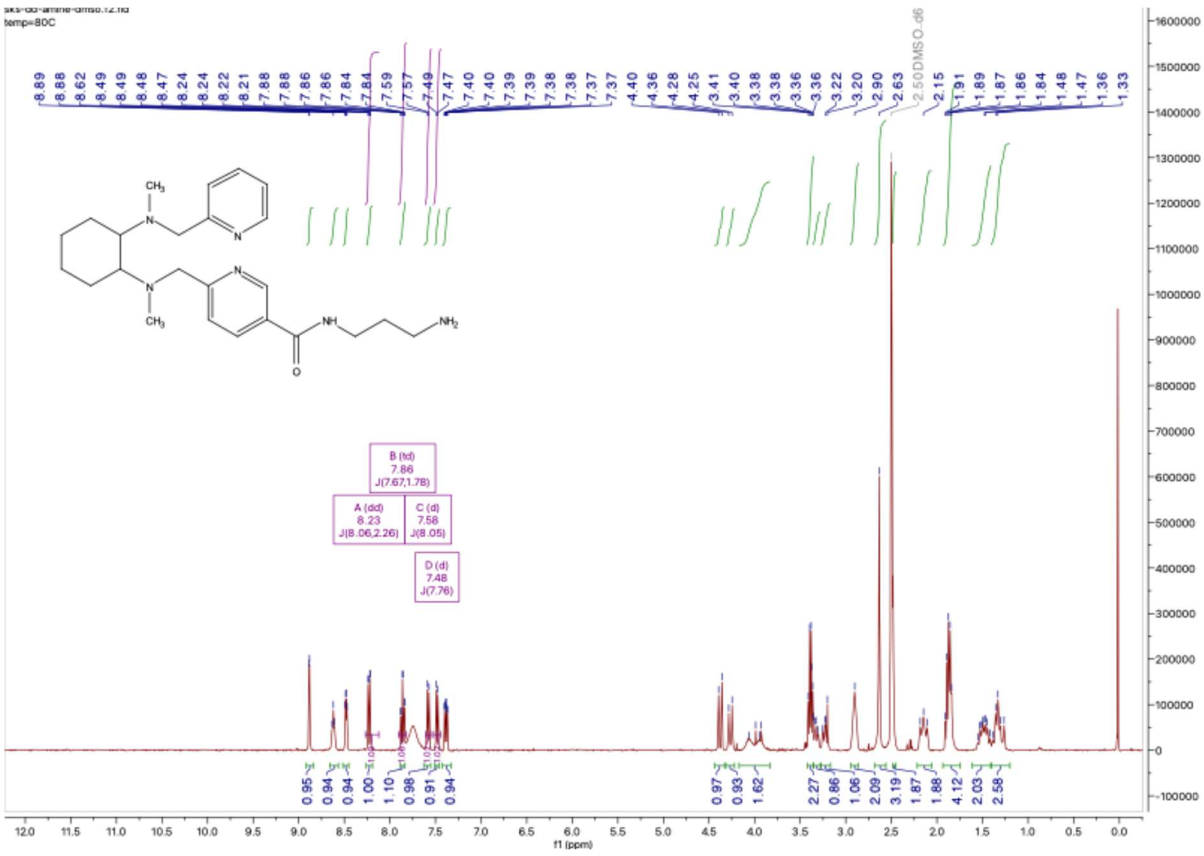

# HR-MS

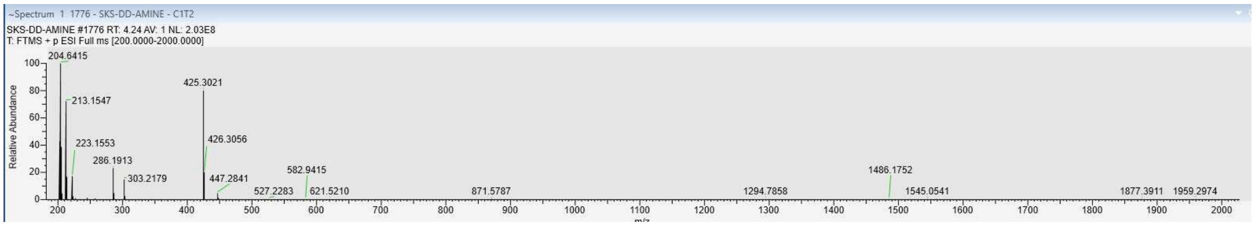
~Spectrum 1 1776 - SKS-DD-AMINE - C1T2
SKS-DD-AMINE #1776 RT: 4.24 AV: 1 NL: 2.03E8
T: FTMS + p ESI Full ms [200.0000-2000.0000]
Relative Abundance
204.6415
213.1547
223.1553
286.1913
303.2179
425.3021
426.3056
447.2841
527.2283
582.9415
621.5210
871.5787
1294.7858
1486.1752
1545.0541
1877.3911
1959.2974

### HPLC:

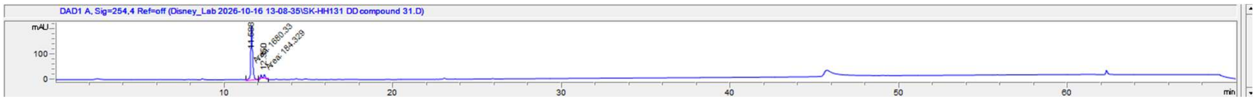
DAD1 A, Sig=254,4 Ref=off (Disney_Lab 2026-10-16 13-08-35\SK-HH131 DD compound 31.D)
mAU
min

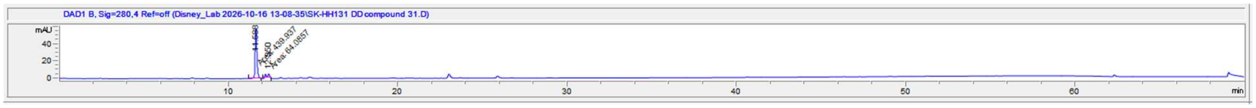
DAD1 B, Sig=280,4 Ref=off (Disney_Lab 2026-10-16 13-08-35\SK-HH131 DD compound 31.D)
mAU
min
